# Mechanical Compression Creates a Quiescent Muscle Stem Cell Niche

**DOI:** 10.1101/2021.10.02.462865

**Authors:** Jiaxiang Tao, Mohammad Ikbal Choudhury, Debonil Maity, Taeki Kim, Sean X. Sun, Chen-Ming Fan

## Abstract

Skeletal muscles can regenerate throughout life time from resident Pax7-expressing (Pax7^+^) muscle stem cells (MuSCs)^1–3^. Pax7^+^ MuSCs are normally quiescent and localized at a niche in which they are attached to the extracellular matrix basally and compressed against the myofiber apically^3–5^. Upon muscle injury, MuSCs lose apical contact with the myofiber and re-enter cell cycle to initiate regeneration. Prior studies on the physical niche of MuSCs focused on basal elasticity^6,7^, and significance of the apical force exerted on MuSCs remains unaddressed. Here we simulate MuSCs’ mechanical environment *in vivo* by applying physical compression to MuSCs’ apical surface. We demonstrate that compression drives activated MuSCs back to a quiescent stem cell state, even when seeded on different basal elasticities. By mathematical modeling and manipulating cell tension, we conclude that low overall tension combined with high edge tension generated by compression lead to MuSC quiescence. We further show that apical compression results in up-regulation of Notch downstream genes, accompanied by increased levels of nuclear Notch. The compression-induced nuclear Notch is ligand-independent, as it does not require the canonical S2-cleavage of Notch by ADAM10/17. Our results fill the knowledge gap on the role of apical tension for MuSC fate. Implications to how stem cell fate and activity are interlocked with the mechanical integrity of its resident tissue are discussed.

---

Skeletal muscles consist of post-mitotic syncytial myofibrils that generate contractile forces for body movement^8^. They have tremendous ability to regenerate after injury mainly owing to the resident Pax7-expressing (Pax7+) muscle stem cells (MuSCs)^1,9^, also known as satellite cells (SCs)^1,4^. Without injury, Pax7+ MuSCs are mostly quiescent. Upon muscle damage, Pax7^+^ MuSCs retain contact with the basement membrane of the dead myofiber (i.e., the ghost fiber^5^), renter cell cycle, become myogenically committed and subsequently differentiate and fuse to form new myofibers^10^. A fraction of proliferative MuSCs self-renew and return to quiescence to maintain stem cell pool as regenerative cycle completes^8^. Molecular and genetic studies have uncovered genes and pathways regulating progressive states of MuSCs during the regenerative cycle^11^. On the other hand, the stiffness of extracellular matrices (ECM) has been shown to play a role in their stemness. For example, laminin-coated hydrogels at an elasticity of 12 KPa, mimicking the physiological elasticity^12^, provide an optimal condition for self-renewal division of MuSCs in culture^7,12–14^. Whereas, culturing them on collagen I fibrils with an elasticity of 2 KPa in conjunction with a synthetic media has been shown to prevent their activation^6^. By contrast, mechanical manipulations at the apical surface of MuSCs have not been conducted to examine the impact on cell fate/state.

Intravital imaging of YFP-marked Pax7^+^ MuSCs revealed their progressive changes of morphology during early injury-regeneration process^5^. In uninjured muscles, MuSCs display a flat and elongated shape: cell dimension perpendicular to myofiber (height) is ~ 4 μm, and cell dimension parallel to myofiber (length) is ~17 μm (Extended Data Fig. 1). At 1-day post injury (1dpi), they are ~5.5 μm in cell height, and ~6 μm in cell length. Whereas at 3dpi, many YFP-marked cells (a collection of MuSCs and progenitors) are proliferating and/or migrating, and their cell height is ~10 μm (with dynamically changing lengths). Changing morphologies of MuSCs have also long been noted on cultured single myofibers^15^, suggesting diverse mechanical tension distributions^16–18^. We were particularly intrigued by the intravital imaging data that the cell shape change at 1dpi occurs prior to MuSC proliferation, and that MuSCs on uninjured myofibers immediately adjacent to an injury site do not appear activated^5^. Thus, we reasoned to examine whether that the compression force exerted on MuSCs by intact myofibers plays a role for MuSC quiescence.

To investigate the link between cellular morphology/mechanical tension and MuSC cell state, we established a system to simulate the mechanical environment for quiescent MuSC. We microfabricated a device that contains a PDMS film with vertical pillars of an average height of 4 μm underneath (Method, Extended Data Fig. 2)^19,20^. MuSCs cultured under this device are presumably compressed to a cell height of 4 μm, mimicking their intact niche. Pax7^+^ MuSCs were isolated from transgenic Pax7-ZSgreen mice^21^ via FACS (Fig. 1a). They were seeded on Matrigel/fibronectin-coated plastic and cultured for two days in growth media. At this time, the cell height was measured at ~8.7 μm using confocal imaging. After adding the compression device, we found that cells were indeed at the expected average height of ~4 μm, which is determined by the confocal imaging (Extended Data Fig. 3b).

**Fig. 1.**
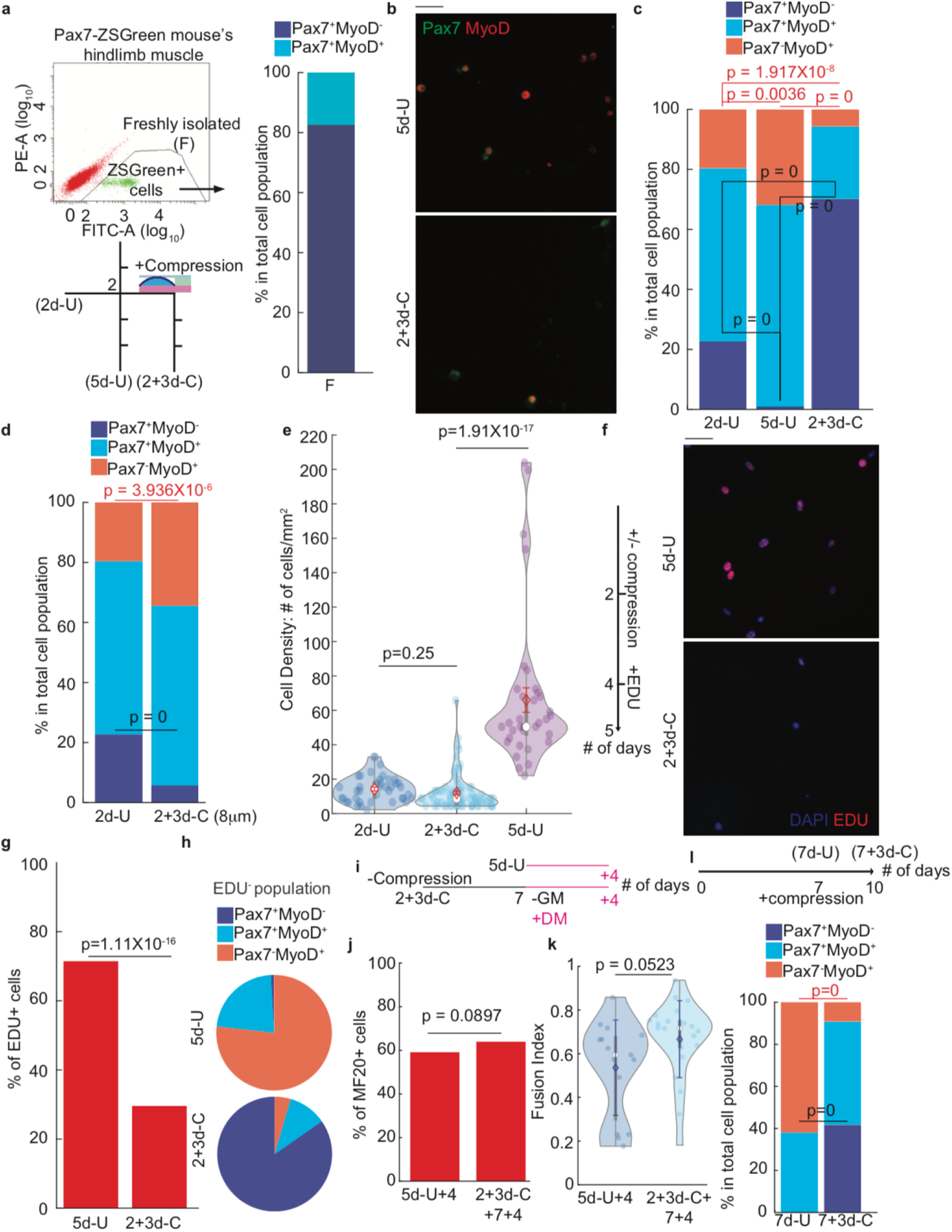
Manipulating cell fate using mechanical compression. **a**, Experimental and FACS setup: Pax7+ was isolated from the hindlimb muscle of Pax7-ZSGreen mice, 3-6 months old. Cytospin result for freshly isolated (F) cells. (Three trails, 741 cells) **b,** Examples of 5d-U and 2+3d-C cells (scale bar 25 μm). **c,** Cell fate evaluation of 2d-U, 5d-U and 2+3d-C cells. (Four trails for 2d-U, 260 cells; Five trails for 5d-U, 3,752 cells; Thirteen trails for 2+3d-C, 420 cells). **d,** Cell fate comparison between 2d-U and 2+3d-C cells with 8μm-high-pillar compression device. (2d-U data is the same with **(c)**; Six trails for 2+3d-C (8μm), 1,032 cells). **e**, Cell densities comparison between 2d-U, 5d-U and 2+3d-C cells. (Same sample as in **(c)** 40 fields of view for 2d-U, 73 fields of view for 2+3d-C, and 37 fields of view for 5d-U). **f,** EDU incorporation setup and examples of 5d-U and 2+3d-C cells (scale bar 25 μm); **g-h,** Fraction of EDU+ cells of 5d-U and 2+3d-C cells **(g),** and cell fate component of EDU-population of 5d-U and 2+3d-C cells (**h)** (6 trails for 5d-U, 3,752 cells; 8 trails for 2+3d-C, 180 cells). **i,** Experimental setup to test the redifferentation capacity of compressed cells. **j-k,** Redifferentation capacity of compressed cells compared to that of 5d-U cells: fraction of MF20+ population **(j),** and fusion index **(k)**. (4 trails for 2+3d-C+4+7; 810 cells; 3 trails for 5d-U+4, 450 cells). **l,** Cell fate comparison between 7d-U and 7+3d-C cells. (3 trails for 7d-U, 1,831 cells; 3 trails for 7+3d-C, 602 cells). Data in **(c), (d), (g), (j)** and **(l)** was presented with overall fraction. *p* value was assessed based on two-tailed Cochran-Mantel-Haenszel test using Matlab. **(e)** and **(k)** data was presented with mean ±s.d. *p* value was assessed with student’s two-tail t-test using Matlab. Comparison was considered significant if *p* ≤ 0.05

To evaluate the role of compression on the cell fate of MuSCs, we examined the expression of Pax7 and MyoD: Pax7^+^MyoD^−^ for stem cells, Pax7^+^MyoD^+^ for progenitors, and Pax7^−^MyoD^+^ for differentiation committed cells^22^. Freshly isolated MuSCs (F) are all Pax7^+^, with some also expressing MyoD (Fig. 1a). After initial two days of culture without compression (2d-U), the majority of cells expressed MyoD (>80%), indicative of MuSC activation. MuSCs were then, either left uncompressed for additional 3 days (5d-U) or subjected to compression for the same period (2+3d-C). The Pax7^+^MyoD^−^ population was almost depleted in the 5d-U. By contrast, 2+3d-C MuSCs contained a significantly larger fraction of Pax7^+^MyoD^−^ cells and smaller fractions of Pax7^+^MyoD^+^ and Pax7^−^MyoD^+^ cells, compared to those of the 5d-U and 2d-U (Fig. 1b, c, Extended Data Fig. 4a-c). Given the ~ 8.7 μm cell height of 2d-U MuSCs, we applied a device with pillars of 8 μm height (for ~ 10% compression) as an additional control and obtained a similar cell fate distribution as the 5d-U condition, albeit with a slight increase of Pax7^+^MyoD^−^ cells (Fig. 1d). We noticed that the cell density of 2+3d-C MuSCs was similar to that before compression (2d-U), and lower than that of the 5d-U (Fig. 1e). Indeed, using EdU-incorporation assay, we found that the 2+3d-C cell population was much less proliferative than the 5d-U population (Fig. 1f,g). Within the EdU^−^ population, the majority of 2+3d-C cells were Pax7^+^MyoD^−^, while the majority of 5d-U cells was Pax7^−^MyoD^+^ (Fig. 1h). Thus, limiting MuSC to 4 um cell height enriches for a non-proliferative Pax7^+^MyoD^−^ state.

Importantly, after removal of the compression pillar, the 2+3d-C cells could increase cell density in ensuing days (Extended data Fig. 4d,e). When they reached a cell density comparable to that of 5d-U cells, we switched them to differentiation medium, and found them to express myosin heavy chain (MF20) and fuse as multinucleated myotubes. The percentage of MF20^+^ cells and the fusion index of de-compressed cells were similar to those of the 5d-U cells subjected to the same differentiation condition (Fig. 1i-k). Together, these data indicate that mechanically compressing MuSCs to 4 μm cell height - equivalent to their physiological cell height in uninjured muscle - can efficiently drive them into a non-proliferative Pax7^+^MyoD^−^ state while retaining the potential for expansion and myogenic differentiation.

We also assessed the effect of compression on MuSCs that have been cultured longer. After 7 days of culture (7d-U), Pax7^−^MyoD^+^ cells were the majority (> 60%), while Pax7^+^MyoD^−^ cells were scarce. After applying compression for additional 3 days (7+3d-C), we found that the Pax7^−^MyoD^+^ fraction decreased substantially (to <10%). Within the Pax7^+^cell fraction, over half was Pax7^+^MyoD^−^ (Fig. 1l). The decrease of the Pax7^−^MyoD^+^ fraction under 2+3d-C and 7+3d-C conditions (from their respective starting points) suggests that some Pax7^−^MyoD^+^ cells may turn on Pax7 under compression.

We next modeled the key change(s) in mechanical tension of cells under compression that drives MuSCs quiescence. A force-balance-based mathematical model^17,18^ is used to estimate the relative change of cellular tension along the two principal directions, lateral (azimuthal) and axial, when the compression is applied (Extended Data Fig. 5). Two assumptions in this model are: 1) cell volume is maintained at a constant at any given cell heights and 2) cell tension is linearly correlated with local cell geometry, i.e., local radius of curvature^17,23^ (Methods). We measured the cell volume of MuSCs and found no significant changes before and immediately after compression, validating the first assumption (Extended Data Fig. 3b). The model predicts that, under compression, a cell experiences a higher axial tension and a lower overall tension, as it becomes flattened and axially extended (Fig. 2a). We calculated that a 4 μm height limitation applied to MuSCs at ~8.7 μm cell height (at day 2 in culture) brought ~55% of lateral compressive strain, resulted in a ~60% reduction of overall tension and a ~2-fold increase in axial tension (Fig. 2a). To visualize these changes, we probed for the phospho-myosin light chain (pMLC) level as a proxy for cell tension^16,24,25^; both the overall intensity and the edge-to-center ratio (for the relative axial tension, Method) of pMLC signals were determined. Compared to 5d-U cells, 2+3d-C cells had a significantly lower overall pMLC level and higher edge-to-center pMLC ratio (Fig. 2b, c, Extended Data Fig. 6a). Thus, experimental data support our modeling that mechanical compression causes lower overall tension and higher axial tension.

**Fig. 2.**
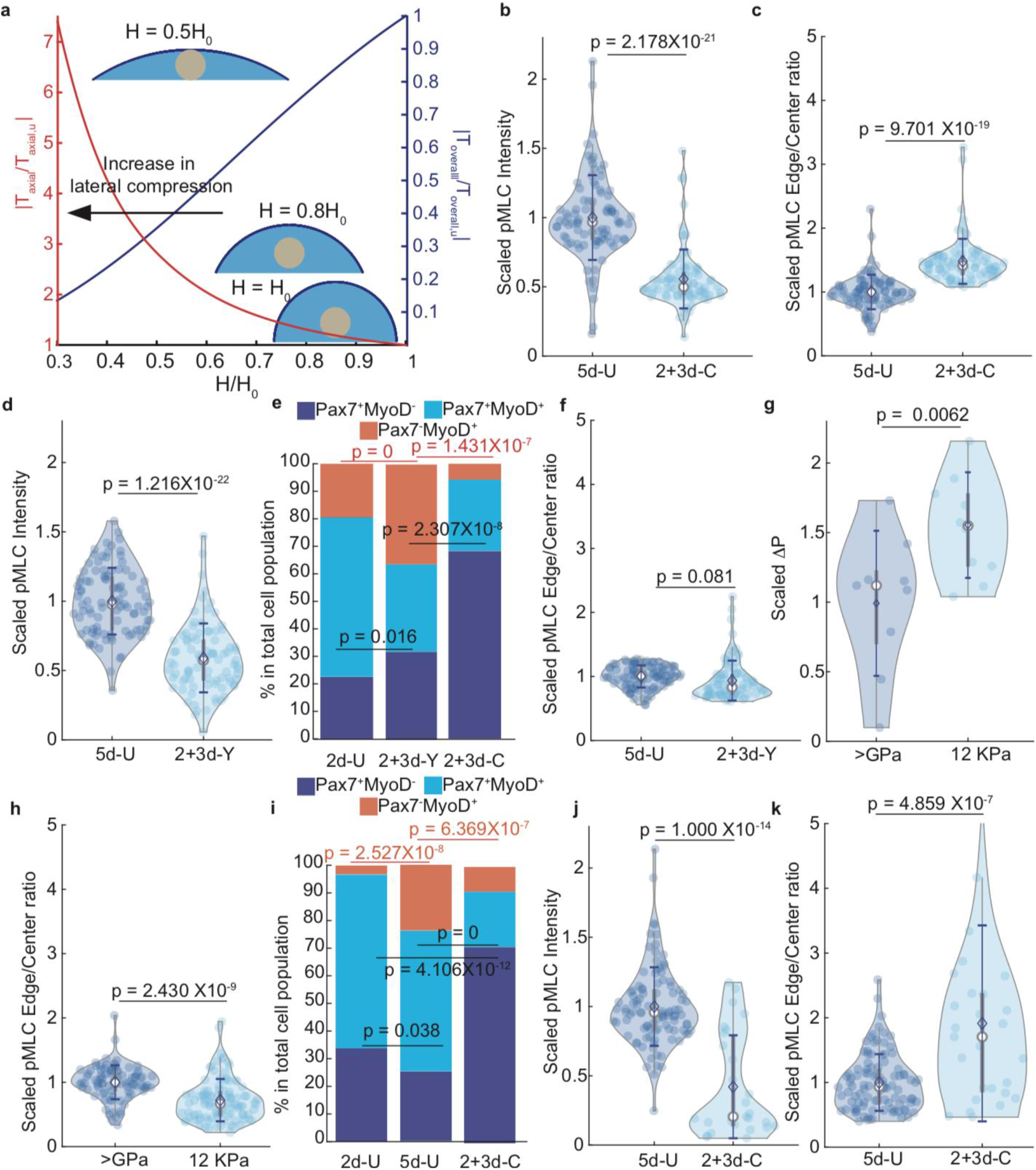
Tension analysis for MuSCs under different conditions. **a,** Mathematical model prediction of change in average tension distribution in terms of compressive strain. **b-c,** pMLC distribution of 5d-U and 2+3d-C cells. This is in terms of overall pMLC **(b)** and pMLC Edge/Center ratio **(c)**. (3 trails were done both 5d-U and 2+3d-C cells. Each trail is scaled with the mean value of its respective 5d-U. 86 cells for 5d-U and 84 cells for 2+3d-C). **d and f,** pMLC distribution of 5d-U and 2+3d-Y cells. This is in terms of overall pMLC **(d)** and pMLC Edge/Center ratio **(f)**. (3 trails were done both 5d-U and 2+3d-Y cells. Each trail is scaled with the mean value of its respective 5d-U. 95 cells for 5d-U and 87 cells for 2+3d-C). **e,** Cell fate comparison between 2d-U, 2+3d-Y and 2+3d-C cell. (2d-U cell fate data is same as in **Fig. 1c**. 4 trails for 2+3d-Y, 268 cells; 3 trails for 2+3d-C cells, 77 cells). **g,** Pressure comparison between 5d-U cells seeded on plastic (stiffness > GPa) and on 12 KPa hydrogel. (Three trails for plastic-seeded cells, 12 cells. Three trails for 12 KPa hydrogel-seeded cells, 10 cells). **h,** pMLC Edge/Center ratio comparison of 5d-U cells seeded on plastic and on 12 KPa hydrogel. (5d-U cells on plastic is same as in **(b)** and **(c)**. Three trails for 5d-U cells seeded on 12 KPa hydrogel, 100 cells). **i**, Cell fate comparison between 2d-U, 5d-U and 2+3d-C cells seeded on 12 KPa hydrogel. (Three trails for 2d-U, 148 cells. Four trails for 5d-U 532 cells, Four trails for 2+3d-C, 228 cells). **j-k,** pMLC distribution of 5d-U and 2+3d-C cells seeded on 12 KPa hydrogel. This is in terms of overall pMLC **(j)** and pMLC Edge/Center ratio **(k)**. (5d-U data is the same as in **(h)**. Each trail is scaled with the mean value of its respective 5d-U. Three trails for 2+3d-C, 27 cells). Modeling results presented in **(a)** is up to second-order accuracy (Method). **(b), (c), (d), (f), (g), (h), (j)** and **(k)** data were presented with mean ±s.d. *p* value was assessed with student’s two-tail t-test using Matlab. Data in **(e)** and **(i)** were presented with overall fraction. *p* value was assessed based on two-tailed Cochran-Mantel-Haenszel test using Matlab. Comparison was considered significant if *p* ≤ 0.05

We next examined how changing cell tension impacts MuSC fate. First, we reduced overall tension by treating the cells with Y-27632 (which inhibits ROCK, an upstream effector of pMLC) after 2 days of culture and assaying them at day 5 (2+3d-Y cells). The 2+3d-Y cells had a lower overall pMLC level as the 2+3d-C cells (50~60% reduction, Fig. 2d), when compared to 5d-U cells. Atomic Force Microscopy (AFM, Method) determined a ~50% reduction in cross-membrane pressure, ΔP, which is linearly scaled with overall tension by the factor of cell height (Method)^18^, of the 2+3d-Y cells relative to that of 5d-U cells (Extended Data Fig. 6b), lending support for pMLC level as a proxy for cell tension. Interestingly, 2+3d-Y cells had only a moderately higher Pax7^+^MyoD^−^ fraction and significantly lower Pax7^+^MyoD^−^ fraction than those of 2+3d-C cells, respectively (Fig. 2e), but a higher Pax7^+^MyoD^−^ fraction than of 5d-U cells (Extended Fig. 6c). Although 2+3d-C and 2+3d-Y cells had a similarly low overall pMLC level, the latter showed no increase of pMLC edge-to-center ratio, relative to 5d-U cells (Fig. 2f, Extended Data Fig. 6a). Thus, lowering overall tension without increasing axial tension only yielded a moderate increase of Pax7^+^MyoD^−^ cell fraction. Applying compression onto Y-27632-treated MuSCs resulted in substantial cell loss, precluding assessment. Next, we increased the overall cell tension by culturing MuSCs on Matrigel/fibronectin-coated 12 KPa hydrogel^16^. AFM measured a ~60% higher ΔP (reflecting an increase in overall cell tension) of MuSCs on 12 KPa, than that of MuSCs on plastic (Fig. 2g). The pMLC edge-to center ratio of cells on 12 KPa was ~10-15% lower than that of cells on plastic (Fig. 2h), which in fact reflects ~30% higher absolute axial tension after taking the increased total tension into account. More MuSCs on the 12 KPa were maintained as Pax7^+^MyoD^−^ in 2d-U and 5d-U settings, compared to those on plastic, respectively (Extended Data Fig. 6d). This is consistent with the report that 12 KPa is better in maintaining self-renewal divisions of MuSCs^7,12^. The Pax7^+^MyoD^−^ fraction were decreased from 2d-U to 5d-U on 12 KPa (Fig. 2i), and considerably less than that of compressed MuSCs on plastic. Thus, simultaneously elevating overall and axial tensions were not as effective in restoring Pax7^+^MyoD^−^ fate as compression. By contrast, after compression on 12 KPa, 2+3d-C cells showed a lower overall level and higher edge-to-center ratio of pMLC, compared to 5d-U cells, following the similar trend of pMLC distribution of 2+3d-C cells on plastic (Fig. 2 j, k, Extended Data Fig. 6e). Importantly, the Pax7^+^MyoD^−^ fractions of 2+3d-C cells, on plastic and 12 KPa, were also similar (Fig. 2i, Extended data Fig. 6d,f). These results together indicate that overall low tension and high axial tension achieved concomitantly by mechanical compression are most effective in supporting the Pax7^+^MyoD^−^ fate. MuSCs within the uninjured muscle do display flattened and extended shape (Extended Data Fig. 1), implying that they experience higher axial tension and lower overall tension as mechanically compressed MuSCs.

To uncover the molecular basis, we interrogated the RNA-seq data among 2+3d-C, 5d-U, and freshly isolated MuSCs (Fig. 3a). Principal component analysis indicates that 2+3d-C cells is more similar to freshly isolated cells, rather than 5d-U cells (Extended Data Fig. 7a). Genes known to be associated with MuSC quiescence^12,26–30^ are upregulated in 2+3d-C cells (Fig. 3b). Whereas, genes associated with MuSC activation, proliferation or myogenic differentiation^26,28^ are upregulated in 5d-U cells (Fig. 3c). Notably, *Pax7*, *calcitonin receptor* (*Calcr*), and *collagen 5* genes (*Col5a1* and *Col5a3*) are upregulated in 2+3d-C cells, even though there exist non-Pax7^+^MyoD^−^ cells. High levels of *Pax7* are linked to MuSC stemness^31^ and *Calcr* and *Col5a1/3* are critical for MuSC quiescence^30,32^. Together with the EdU data (Fig.1h), 2+3d-C Pax7^+^MyoD^−^ cells represent a cell fate/state mimicking quiescent MuSCs^28,29^.

**Fig. 3.**
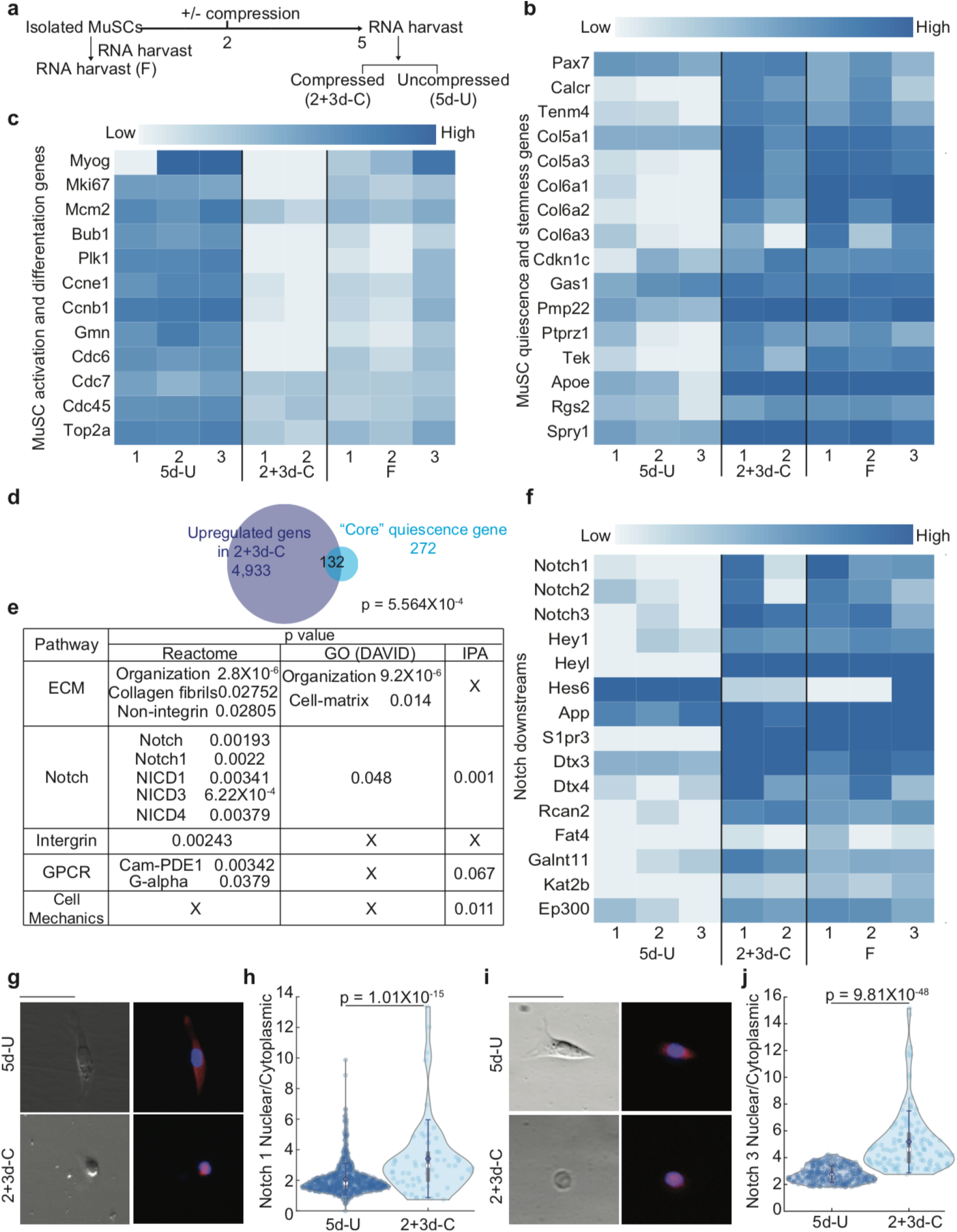
Transcriptome comparison between 2+3d-C, 5d-U and freshly isolated (F) cells. **a,** Experimental setup. **b,c,f,** FPKM readings of MuSCs’ quiescence and stemness **(b)**, activation and differentiation **(c)**, and Notch downstream genes **(f)**. **d**, Comparison between upregulated genes in 2+3d-C cells and “core” quiescence genes. **e,** Pathway analyses on overlapped “core” quiescence genes. **g-j,** Notch1 **(g,h)** and 3 **(i,j)** nuclear/cytoplasmic ratio comparison between 5d-U and 2+3d-C with respective examples **(g** and **i)**(Scale bar 25 μm). (**(h):** Four trails for 5d-U (+DMSO), 685 cells. Four trails for 2+3d-C (+DMSO), 61 cells. **(j)** Five trails for 5d-U (+DMSO), 291 cells. Three trails for 2+3d-C (+DMSO), 124 cells). **b,c,f** Raw reads number for each gene is generated by R, and FPKM reading, as well as respective *p* value are calculated by Matlab’s Genomic package All presented genes have *p* value smaller than 0.05. *p* value for overlapping genes in **(d)** was calculated by Fisher Extract method, and by each respective pipeline in **(e)** If *p>*0.1 for that specific pipeline, “X” is marked. **(h,j)** data were presented with mean ±s.d. *p* value was assessed with student’s two-tail t-test using Matlab. Comparison was considered significant if *p* ≤ 0.05.

Recent studies comparing steady state and nascent transcriptomes of freshly isolated MuSC to those fixed *in situ* revealed substantial differences^28,29^. Genes up-regulated in fixed MuSCs are not identical between steady state and nascent transcriptomes, but do show overlap of 272 genes. We used these genes as core quiescent genes for pathway analysis (Extended data Fig. 7b), and found an enrichment for Notch, Hedgehog, and G-protein coupled receptor/cyclic-AMP signaling using the reactome tool; Pathway enrichment by GO (DAVID) and IPA were also performed (Extended Data Fig. 7c). In contrast, the up-regulated genes in 2+3d-C MuSCs show pathway enrichment for Notch signaling, as well as actin-related cell mechanics and cell-ECM adhesions (likely due to compression; Extended Data Fig. 7d). We then compared the 272 core quiescent genes to the up-regulated genes in 2+3d-C MuSCs (Fig. 3d), and the 132 overlapped genes (*p* = 5.564 × 10^−4^, Fisher Exact) were subjected to the same analyses. Notch signaling is still enriched in all three pipelines (Fig. 3e). Notch1-3 have been implicated in regulating MuSCs: Notch1 and 2 play a concerted role in maintaining their quiescence and stemness, whereas Notch3, their return to quiescence^33,34^. Additionally, direct downstream genes of Notch, such as *Her* genes^26,30,35^, are up-regulated in 2+3d-C cells (Fig. 3f). Thus, comporessed MuSCs show gene signatures for Notch signaling.

To demonstrate Notch activation by compression, we examined the cellular localization of Notch1 and Notch3 in MuSCs. Notch activation requires binding and endocytosis of its ligand Delta (Dll, or Dll-related members in mammals) by neighboring cells to generate a pulling force necessary for a conformation change of the juxta-membrane NRR domain of Notch to allow S2-cleavage by ADAM10 and 17, followed by an intramembrane S3-cleavage by γ-secretase^36^. S3-cleaved Notch intracellular domain then enters the nucleus to affect gene expression^36^. Indeed, compared to 5d-U cells, 2+3d-C cells displayed nuclear enrichment of Notch1 and Notch3 (Fig. 3g-j), providing evidence for Notch pathway activation.

As we did not provide exogeneous Dll with a pulling force, it is unlikely that nuclear-enriched Notch1 and Notch3 are mediated by the canonical mode. In addition, 2+3d-C MuSCs are singly isolated and mostly non-proliferative, making a low probability for paracrine Dll-Notch signaling via cell-cell contact^37^. We therefore considered a Dll-independent cell-autonomous mode of Notch activation by compression. If so, S2-cleavage may not be needed, while the S3-cleavage should still be required for nuclear Notch. To test this, we inhibited S2-cleavage by blocking ADAM10 and 17 with GI254023X and TAPI0, respectively, and S3-cleaveage by blocking γ -secretase with DAPT. Notch3 nuclear enrichment and cell fate evaluation of 5d-U and 2+3d-C MuSCs under these pharmacological treatments were compared to those of mock-treated controls (Fig. 4a; Method). Inhibition of either S2- or S3-cleavage increased the Pax7^−^MyoD^+^ cell fraction and decreased nuclear Notch3 to a similar level in 5d-U MuSCs (Fig. 4b, c). Compression-induced increases of Pax7^+^MyoD^−^ cell fraction and nuclear Notch3 were only abolished by inhibiting S3-cleavage (Fig. 4d, e, Extended Data Fig. 8). Given the similar results of blocking S2 or S3-cleavage were observed for 5d-U cells, the difference of blocking them found in 2+3d-C cells implies that the canonical Dll-dependent S2-cleavage of Notch is not critical for MuSC stemness under compression.

**Fig. 4.**
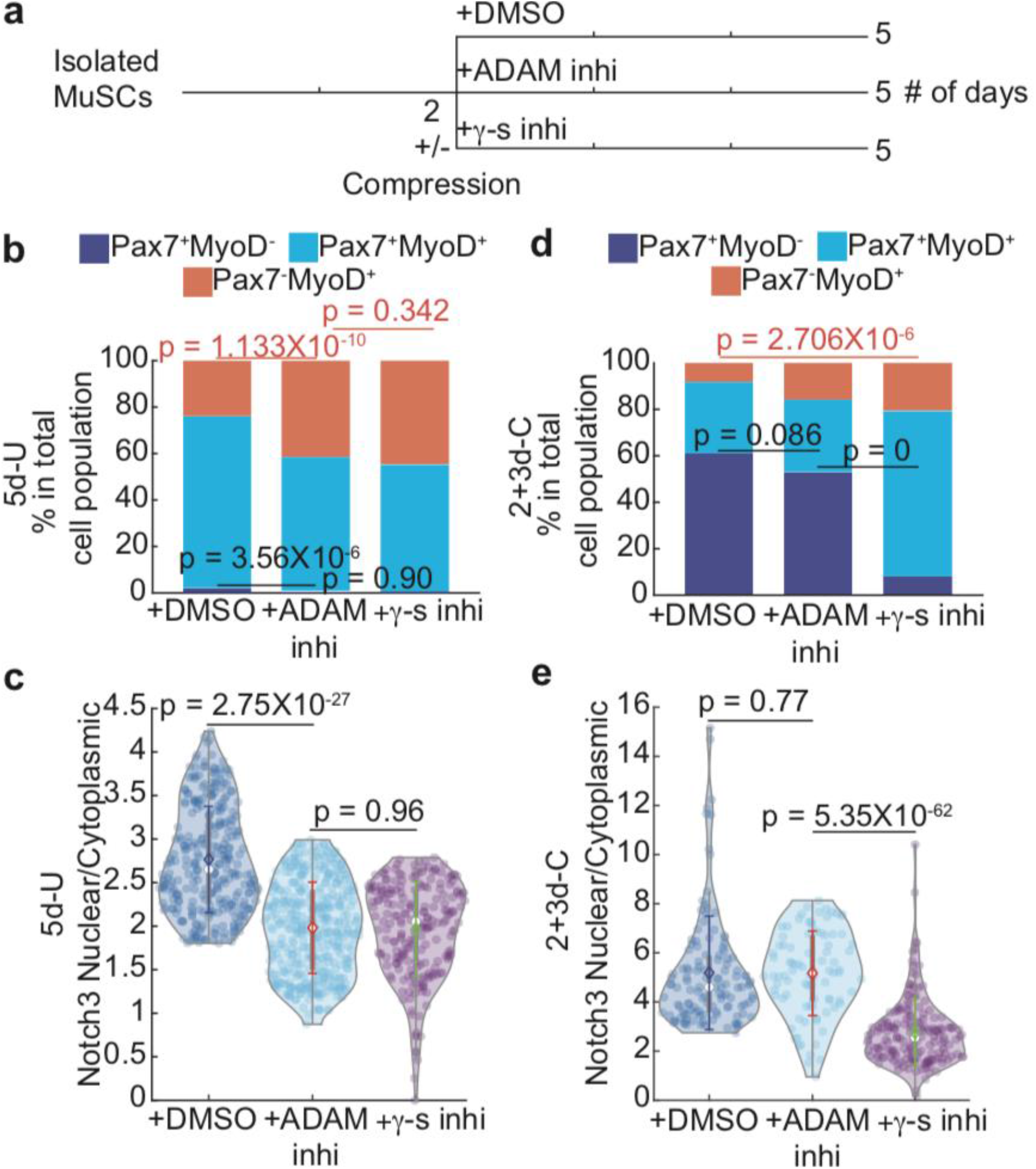
Notch nuclear enrichment assay and cell fate evaluation after blocking S2 (+ADAM inhi) or S3 (+γ-s inhi). **a,** Experimental setup. **b-c,** 5d-U cells’ fate (**b)** and Notch3 nuclear/cytoplasmic ratio **(c)** comparison after blocking S2 or S3. (**(b):** Seven trails for control (+DMSO), 1,303 cells; three trails for +ADAM inhi, 332 cells; Three trails for +γ-s inhi: 596 cells. **(c):** control (+DMSO) data is same as in **Fig. 3j.** Four trails for +ADAM inhi, 423 cells. Three trails for +γ-s inhi, 215 cells). **d-e,** 2+3d-C cells’ fate (**d)** and Notch3 nuclear/cytoplasmic ratio **(e)** comparison after blocking S2 or S3. (**(d):** Seven trails for control (+DMSO), 280 cells; Five trails for +ADAM inhi, 183 cells; Four trails for +γ-s inhi: 715 cells. **(e):** control (+DMSO) data is same as in **Fig. 3j.** Three trails for +ADAM inhi, 88 cells. Three trails for +γ-s inhi, 169 cells). Data in **(b)** and **(d)** was presented with overall fraction. *p* value was assessed based on two-tailed Cochran-Mantel-Haenszel test using Matlab. **(c)** and **(e)** data was presented with mean ±s.d. *p* value was assessed with student’s two-tail t-test using Matlab. Comparison was considered significant if *p* ≤ 0.05

Other than aberrant activation by cancer promoting mutations in Notch(es)^38^, Dll-independent physiological activation of Notch has been noted, particularly during T-cell development^39^. Cell-autonomous Notch activation is prevented by keeping Notch levels on the plasma membrane in check via endo-lysosomal degradation machinery^40,41^. Conditions that alter the trafficking dynamics can lead to intracellular Notch activation. It is thought that the S2-cleavage is still needed to expose a cleaved peptide stump for recognition by γ-secretase in endo-lysosomal vesicles^40^, and ADAM 17 is suggested to mediate ligand-independent S2-cleavage^40,42^. By contrast, compression-induced nuclear Notch enrichment occurred when we inhibited both ADAM10 and ADAM17. While we cannot exclude the possibility that another ADAM-related protease(s) mediates compression-induced Notch activation, we propose a mechanical mechanism in which the combined low overall and high relative axial tensions by compression create a membrane topology permissible for eventual S3-cleavage of Notch, distinct from existing proposals for Dll-independent mechanisms^40,41,43^.

## Conclusion

The concept of apically compressed physical micro-environment as a quiescent MuSC niche helps explain their activity in vivo. During the injury-regeneration cycle, MuSCs residing over a damaged myofiber segment lose apical contact and become activated for repair/regeneration^5^. The timing of them returning to quiescence must be coordinated with the completion of muscle repair/regeneration to attain proportional regeneration, and the compression force of fully regenerated myofibers can serve as timing control. A similar scenario likely applies to the establishment of the initial quiescent MuSC pool during postnatal muscle growth as myofibers grow to a mature size and exert sufficient compression forces to instruct quiescence. This principle may be extended to the scaling between MuSCs and muscles during adaptive changes of muscle size, such as exercise and aging. Importantly, our in vitro engineered niche activates Notch signaling utilized for MuSC quiescence in vivo. Synthesizing the known signaling pathways and the findings herein, we re-interpret that Cadherins act to decrease overall tension by mediating apical adhesion with the myofiber^44^, Wnt-Rho signaling causes tension re-distribution^44,45^ (Extended data Fig7b.). Both measures can be accomplished by compression to activate Notch. Further, myofiber-expressed Dll has been implicated in MuSC quiescence^46^, and is localized to MuSCs’ cell edge, a region with high axial tension by compression. We therefore propose that Dll-induced and Dll-independent tension-sensing Notch signaling act in parallel to doubly assure MuSC quiescence. Here we addressed a previously unexplored compressive force on stem cell quiescence, contrasting to basal elasticity, stretch force, osmotic pressure, and other mechanical conduits that stimulate stem cell proliferation or differentiation^6,7,47,48^. The rules for stem cell mechanobiology should vary in different tissues, pending on the shape, size, and organization of cell types and ECM therein. It stands to reason that both quiescent and stimulatory niche mechanics in a micro-scale imposed by a resident tissue on a macroscale, are sensed by stem cells to maintain tissue homeostasis in normal physiology, regeneration, and possibly pathological conditions.

## Acknowledgements

We thank Dr. Frederick Tan for training in transcriptome analysis. We also thank Allison Pinder and Dr. Mahmud Siddiqui for technical assistance, and Dr. Liangji Li & Dr. Nathalie Gerassimov for assistance with FACS. C.-M.F. is supported by NIH (grants R01AR060042, R01AR071976 and R01AR072644) and the Carnegie Institution for Science. S.X.S is supported by NIH grant (R01GM134542)

## Contributions

J. T. and C.-M.F. conceived and designed the study and wrote the manuscript. J. T. carried out all the experiments and epifluorescence imaging, except for AFM, which was carried out by M.I.C. J.T. carried out confocal imaging with assistance by M.I.C. D. M. manufactured the compression device. J. T. carried out all the data analysis, with assistance by T.K. J. T. and T. K. carried out mathematical modeling of cell shape, and method of fitting for AFM data. T. K. carried out AFM data fitting. J. T. carried out all the transcriptome analyses. S. X. S provides facility supports.

## Author Approvals

All authors have seen and approved the manuscript. This manuscript hasn’t been accepted published

## Data Availability

All data is available from authors upon reasonable request.

