## Supplemental Methods and Extended Data Figures & Legends for "Mechanical Compression Creates a Quiescent Muscle Stem Cell Niche"

**Methods:**

**Manufacturing Microfluidic Compression Device**

The compression device is fabricated using a soft-lithography technique[49]. The general process is to etch a silicon wafer with UV light with a specific pattern and depth, determined by a pre-designed photomask and a pre-coated photoresists mixture, respectively. The etched wafer is later used as a mold to "shape" the pattern into a soft PDMS film.

First, the compression pillar pattern is designed by AutoCAD (Extended Data Fig. 2) and printed onto a photomask. Each pattern group has a designed size comparable to the area in a well of a 24-well dish (diameter around 15.6 mm), and one photomask contains 8 of such a pattern group. This photomask allows the UV light to etch only at the designated pillar region (Extended Data Fig. 2), creating a depression which shapes the pillar pattern onto a silicon wafer. The silicon wafer is spun-coated with SU8 3005 photoresists (at 3,000 rpm for 1 min) at specific mixing ratio, which allows the UV to etch only ~4 μm into the wafer. We then align the photomask with a silicon wafer using EV620 Double Sided Mask Aligner and expose it to UV.

After the mold has been created following the UV exposure, we spin-coat a thin layer of PDMS (at 100 rpm for 1.5 min) onto the mold and bake the mold at 70 $^{0}$C oven until the PDMS is completely cured. Therefore, this PDMS film contains protruded-out pillars, with the height of ~4 μm, following the caved-in pillar pattern on the mold (referred to as pillar-side) (Extended Data Fig. 2). Then, the PDMS is then carefully peeled off from the mold and attached to a cover glass at the non-pillar side.

### Applying Mechanical Compression to cultured satellite cells

When the compression needs to be applied to cultured cells in a 24-well plate, each pillar group is carefully cut off from the cover glass and directly added to the cell culture with the pillar side facing the cell. Heavy weight is applied at the top (non-pillar side) of the PDMS film to prevent it from floating. After each experiment, each pillar group is taken out of the dish, rinsed with 70% alcohol solution, and is preserved in DI water. To minimize the experimental error brought by the erosion from the pillar, we replace the compression pillars every 10 months.

After application of the PDMS film, the cells are confined into a wide channel (200 μm by 200 μm) between the neighboring pillars and their heights are limited to ~4 μm. To examine this, we measured the actual 2d-U Pax7-ZSGreen cell height using confocal microscopy after adding the compression pillar and found that the average cell height is ~4μm, matching our designed pillar height (Extended Data Fig. 3a,b).

### Mathematical Model Description

The change in local cellular curvature and apical tension due to externally applied mechanical compression can be recapitulated by using a force-balance based mathematical model[18]:

$$(\nabla \cdot \mathbf{n})^{-1} = \frac{T}{\Delta P} \quad (1)$$

in which $\mathbf{n}$ is local unit normal vector. $\nabla$ is operating at local cell membrane surface. $T$ is overall cell tension, and $\Delta P$ is intracellular-to-extracellular hydrostatic pressure difference. By geometric definition, $(\nabla \cdot \mathbf{n})$ equals to the trace of curvature tensor, which describes the local shape of the cell.

To calculate the mean curvature, we parametrize the cell surface using cell local radius, *R*, azimuthal angle $\theta$, and polar angle, $\phi$, as shown Extended Data Fig. 5. If we assume the cell shape is axisymmetric, *R* only depends on azimuthal angle, $\theta$. Therefore, we can write down the location for a given point on the cell surface, using our parametrization:

$$\boldsymbol{x} = (R(\theta)\sin\theta\cos\phi\,, R(\theta)\sin\theta\sin\phi\,, R(\theta)\cos\theta) \quad (2)$$

Therefore, the two tangential vectors, along the azimuthal direction and axial (polar) directions, respectively (Extended Data Fig. 5), are:

$$\frac{\partial \boldsymbol{x}}{\partial \theta} = (R'\sin\theta\cos\phi + R\cos\theta\cos\phi\,, R'\sin\theta\sin\phi + R\cos\theta\sin\phi\,, R'\cos\theta - R\sin\theta)$$

$$\frac{\partial \boldsymbol{x}}{\partial \phi} = (-R\sin\theta\sin\phi\,, R\sin\theta\cos\phi\,, 0) \ (3)$$

in which we define: $(\dots)' = \frac{\partial(\dots)}{\partial\theta}$. Thus, the geometric tensor for the cell is $\boldsymbol{g_{ij}} = \partial_i \boldsymbol{x} \cdot \partial_j \boldsymbol{x}$, which is:

$$\mathbf{g} = \begin{bmatrix} \frac{\partial \boldsymbol{x}}{\partial \theta} \cdot \frac{\partial \boldsymbol{x}}{\partial \theta} & \frac{\partial \boldsymbol{x}}{\partial \theta} \cdot \frac{\partial \boldsymbol{x}}{\partial \phi} \\ \frac{\partial \boldsymbol{x}}{\partial \phi} \cdot \frac{\partial \boldsymbol{x}}{\partial \theta} & \frac{\partial \boldsymbol{x}}{\partial \phi} \cdot \frac{\partial \boldsymbol{x}}{\partial \phi} \end{bmatrix} = \begin{bmatrix} R^2 + R'^2 & 0 \\ 0 & R^2 \sin^2\theta \end{bmatrix} \quad (4)$$

The zero-off-diagonal components of **g** indicates the parametrization described above (Extended Data Fig. 5) is along the principal direction of the surface, which also results curvature tensor with zero-off-diagonal components. Therefore, the diagonal components of curvature tensor, $c_1$ and $c_2$, represent the cell shape along the azimuthal and axial direction, respectively. The local unit normal vector can be calculated based on equation (3): $\boldsymbol{n} = \frac{\frac{\partial x}{\partial \theta} \times \frac{\partial x}{\partial \phi}}{\det(\boldsymbol{g})}$.

To calculate the curvilinear divergent for the local unit normal, we need to first calculate the local curvature tensor: $\Theta = \partial_i \partial_j \boldsymbol{x}_k \boldsymbol{n}_k$, which also has zero off-diagonal components:

$$\Theta = \begin{bmatrix} \frac{\partial^2 \boldsymbol{x}(1)}{\partial \theta^2} & \frac{\partial^2 \boldsymbol{x}(1)}{\partial \theta \partial \phi} \\ \frac{\partial^2 \boldsymbol{x}(1)}{\partial \phi \partial \theta} & \frac{\partial^2 \boldsymbol{x}(1)}{\partial \phi^2} \end{bmatrix} \boldsymbol{n}(1) + \begin{bmatrix} \frac{\partial^2 \boldsymbol{x}(2)}{\partial \theta^2} & \frac{\partial^2 \boldsymbol{x}(2)}{\partial \theta \partial \phi} \\ \frac{\partial^2 \boldsymbol{x}(2)}{\partial \phi \partial \theta} & \frac{\partial^2 \boldsymbol{x}(2)}{\partial \phi^2} \end{bmatrix} \boldsymbol{n}(2) + \begin{bmatrix} \frac{\partial^2 \boldsymbol{x}(3)}{\partial \theta^2} & \frac{\partial^2 \boldsymbol{x}(3)}{\partial \theta \partial \phi} \\ \frac{\partial^2 \boldsymbol{x}(3)}{\partial \phi \partial \theta} & \frac{\partial^2 \boldsymbol{x}(3)}{\partial \phi^2} \end{bmatrix} \boldsymbol{n}(3) \quad (4)$$

Therefore, curvature along the azimuthal direction is then: $c_1 = \mathbf{g}_{\mathbf{11}}^{-\mathbf{1}} \Theta_{11}$, and the curvature along the axial (polar) direction is $c_2 = \mathbf{g}_{\mathbf{22}}^{-\mathbf{1}} \Theta_{22}$. We then substitute: $(\nabla \cdot \mathbf{n}) = \boldsymbol{g}_{11}^{-1} \Theta_{11} + \boldsymbol{g}_{22}^{-1} \Theta_{22}$ back to Eq. (1), we have:

$$(\nabla \cdot \mathbf{n})^{-1} = \left( \frac{2R'^2 + R^2 - RR''}{(R^2 + R'^2)^{1.5}} + \frac{1 - \frac{R'}{R} \cot \theta}{\sqrt{R^2 + R'^2}} \right)^{-1} = \frac{T}{\Delta P} \quad (5)$$

Two boundary conditions are required to solve this equation. First, under compression, the cell height is fixed at $H$: $R(\theta = 0) = H$. Second, based on the symmetric condition: $\frac{dR}{d\theta}(\theta = 0) = 0$. This force balance condition is not sufficient to solve for cellular shape of given cellular height, for the overall cell tension is not known. Therefore, we close the system by assuming: $T = k\left(\frac{1}{c1} + \frac{1}{c2}\right)$ and the cell retains a constant volume, $V_0$, right when the vertical compression is applied (in which $V_0$ is uncompressed cell volume). The constant volume assumption comes from our measurement, shown in Extended Data Fig. 3b. We assume the uncompressed cell has a hemisphere shape, with radius of $R_0$. Then, $V_0 = \frac{2}{3}\pi R_0^3$. Based on the measurement shown in Extended Data Fig. 3a in the main text, $R_0$~ 8.7 μm. The compressed cell volume is then:

$$V = \pi \int_0^{\frac{\pi}{2}} R^2 \sin^2 \theta \, (R \sin\theta - R' \cos\theta) d\theta = V_0 \ (6)$$

**Numerical Results:**

Numerically, we discretized $R(\theta)$ with finite angle increment $d\theta = \frac{\pi}{1000}$. We expand the expression for *R* using Taylor series and approximate the first and second derivative to the second-order accuracy:

$$R'(\theta) = \frac{R(\theta + d\theta) - R(\theta - d\theta)}{2d\theta} + O(d\theta^2)$$

$$R''(\theta) = \frac{R(\theta + d\theta) + R(\theta - d\theta) - 2R(\theta)}{d\theta^2} + O(d\theta^2) \ (7)$$

We then substitute Eq. (7) back to Eq. (5) to obtain discretized force balance equation. Then, we run the iteration to solve for *R* for all discretized $\theta$ with a guessed parameter $\frac{T}{\Delta P}$. We then adjust our guess for $\frac{T}{\Delta P}$ until the computed shape will result a volume, *V* (Eq. 6), is within 1% error of uncompressed cell volume $V_0$. or: $\left|\frac{V - V_0}{V_0}\right| \le 0.01$.

This model predicts both the cell shape and the overall local mechanical tension at a given cell height, *H*. We substituted cell tension *T*, with local cell curvature along the azimuthal and axial (polar) direction using elastic theory: $T \propto c_1^{-1}, c_2^{-1}$. As shown in Fig. 2a in the main text, as compression strain increases, the azimuthal tension reduces while axial (polar) tension increases, meaning that the vertical compression received by the cell from the top brings axial extension.

Based on our measurements of cell shape (Fig. 2a), adding a ~4 μm-pillar to the cell culture applies a ~50% of vertical compression strain to the cell. Given that there is no sudden cell volume change following the vertical compression, the axial tension of the cell increases by ~2 times.

**Atomic Force Microscopy (AFM)**

AFM experiments were done with a Silicon Nitride (SiN) cantilever and tips with a nominal spring constant of 0.01 N/m (Bruker, USA) on an MFP3D (Asylum Research, USA) instrument. Thermal fluctuation method was used to calibrate the stiffness of the cantilever prior to every experiment[50]. To measure the apical stiffness, cells were indented using contact mode, to get force-displacement curves (Extended Data Fig.9a-c). The cell-to-cantilever contact area is estimated to be 1 $\mu m^2$. All data processing was done using Igor pro software (Wavemetrics, USA). The indentation was done by indenting the cells with depth no more than 1.5 μm, so that it would fall into the category of "small indentation", as described in the next section. The indentation increment was set so that there was a total of 400 indentation steps for the depth-force curve for each cell.

**Approximating overall cell tension from AFM data**

Because of the small-strain indentation (< 10% of total cell height), and small contact area between AFM cantilever and the cell surface, the cell remained a constant volume and relatively constant intracellular-to-extracellular pressure difference. The cell membrane surface was parametrized according to the description of Method: **Mathematical Model Description** (Extended Data Fig 5). The mechanical energy function, *E*, described the cellular shape change in response to extracellular force. While *E* is a complicated function, the functional derivative of *E* in terms of cell geometry, *R*, is the classic normal force balance, described in Method: Cell Shape Theory section:

$$\frac{\delta E}{\delta R} = \frac{\Delta P}{\nabla \cdot n} - T \ (1)$$

in which $\Delta P$ is the intracellular-to-extracellular hydrostatic pressure difference, and $T$ is the cortical tension; **n** is the local normal unit vector, and $\nabla \cdot n$ equals to the trace of the curvature tensor at that point. According to the derivation from the Method: Cell Shape Theory section:

$$\nabla \cdot n = \frac{2R'^2 + R^2 - RR''}{(R^2 + R'^2)^{1.5}} + \frac{1 - \frac{R'}{R} cot\ cot\ \theta}{\sqrt{R^2 + R'^2}} \ (2)$$

Assuming the un-indented cell shape is close to a hemisphere with cell height and basal radius of $r_0$ (Extended Data Fig. 5), and the cell remained a relatively constant adhesion radius during the small-strain indentation, Eq. (1) yielded two boundary conditions: $R = r_0$ at $\theta = \frac{\pi}{2}$, which satisfied the small-strain indentation assumption, and $R = r_0 - H$ at $\theta = 0$, in which $H$ is the indentation depth. Since small-strain indentation did not bring significant volume change as well, solving cell tension is to minimize function $E$ in terms of cell geometry (by setting Eq. (1) equals to zero), under the constrain of: $V = V_0 = \frac{2}{3}\pi r_0^3$.

To satisfy this constraint, Eq. (1) was minimized by adding a Lagrangian multiplier term: $E = E + \lambda(V_0 - V)$. The physical meaning of $\lambda$ is the volumetric force which modifies the overall intracellular-to-extracellular pressure so that the cell can keep a constant volume during the geometric changes. Therefore, minimizing Eq. (1) has become:

$$\frac{\delta E}{\delta R} - \lambda \frac{\delta V}{\delta R} = 0 \ (3)$$

$\frac{\delta E}{\delta r}$ is from Eq. (1). Then, couple Eq. (3) equation with:

$$V = \int_0^{\frac{\pi}{2}} R^2 \sin^2\theta \,(R\sin\theta - R'\cos\theta)d\theta = \frac{2\pi}{3} r_0^3 \ (4)$$

Therefore, the functional derivative of *V* in terms of cell geometry, *R*, is: $\frac{\delta V}{\delta r} = \pi(3R^2 \sin^3\theta \ + 2RR'\theta\cos\theta\sin^2\theta\ )$. Substituting this and Eq. (2) to Eq. (3), and couple Eq. (3) with Eq. (4), the cell shape, as well as the volumetric force, $\lambda$, can be solved with given post-indentation cell height *H*

Then, by taking a force balance just at the cantilever-to-cell membrane contact area, the feedback force per unit area from the cantilever is:

$$F = \left(\Delta P + \frac{\lambda V}{A}\right) - T\ (\nabla \cdot n)|_{\theta=\frac{\pi}{2}} \ (5)$$

in which $V$ is the cell volume. $\lambda V$ gives the total forces that are added to the whole cell membrane to maintain a constant volume, and $A$ is the apical area. We adjusted the pressure difference with the added volume-constrain force: $\Delta P = \Delta P + \frac{\lambda V}{A}$

To numerically solve this set of equations, we expand the geometry of the cell using Fourier Series:

$$r = r_0 + \sum_n a_n \cos(n\theta) + \sum_n b_n \sin(n\theta) \quad (6)$$

Considering of boundary conditions discussed above, we have: $\sum_n a_n \cos\left(\frac{n\pi}{2}\right) + \sum_n b_n \sin\left(\frac{n\pi}{2}\right) = 0$, which satisfies the adhesion boundary condition; and $\sum_n a_n = H - r_0$, which satisfies the constant cell height. The coefficients of this expansion should give a cell geometry that satisfies Eq. (4) and $\frac{d\lambda}{d\theta} = 0$, which is derived from Eq. 3. $R$ is expanded up to the second term of the Fourier series, which gives 4 parameters.

With the best fit, we calculated the theoretical $\frac{F}{\Delta P}$ in terms of the indentation strain (Extended Data Fig. 9 d-f, green line). This means, regardless of cell type and mechanical condition, any indentation strain-stress curve should collapse with this theoretical indentation strain-$\frac{F}{\Delta P}$ curve by timing it with a constant. This constant is the estimated $\frac{1}{\Delta P}$ for that cell. Examples shown in Extended Data Fig. 9d-f show that indeed we can fit the indentation strain-stress curve to the theoretical curve for cells under three different conditions (i.e., 5d-U cells on plastic and 12 KPa and 2+3d-Y cells on plastic). We can then estimate the overall tension from the fitted pressure: $T_{overall} \sim \Delta P$ L. L is the typical length scale of the cell, usually in the order of the un-indented cell height, $r_0$. Even though cell heights between 5d-U cells on plastic or on 12 KPa or 2+3d-Y cells may be different, they are in similar order of magnitude. Therefore, using fitted $\Delta P$ to approximate overall tension reflects the general trend of overall tension between these three conditions. Specifically, regardless of whether the cells are seeded on 12 KPa hydrogel or plastic, a 4 mm compression decreases the overall pMLC level by 50% (Fig. 2). This indicates that the cellular height of plastic seeded cells is similar compared to that of 12 KPa hydrogel-seeded cells.

**Immunofluorescence and Imaging**

Cells were fixed by 4% paraformaldehyde (PFA) in phosphate-buffered saline (PBS) in 24-well plate dish for 10 minutes at room temperature. They were then permeabilized by 0.5% Triton in PBS for 10 minutes and then blocked in 100% Normal Goat Serum (NGS, Blocking Buffer) for 1 hour at room temperature. Primary antibodies were diluted in the blocking buffer at specified concentrations listed below and applied to the samples at each well, which were rocked at 4 $^{0}$C overnight. Following day, samples were washed twice with 0.1% Triton in PBS, treated with 4’,6-diamidino-2-phenylindole (DAPI, 1:1000 dilution) plus appropriate secondary antibodies in blocking buffer, and were rocked at room temperature for 1 hour, protected from light. They were then washed three times with 0.1% Triton in PBS prior to imaging. The negative control samples for each 24-well plate were subjected to the same procedure in parallel without primary antibodies to the specific antigen, in order to determine the cut-off intensity values for positive signal to that antigen.

**Epi-fluorescent Imaging**

Fluorescent images were taken by a HAMAMATSU camera (Model C13440) mounted on a Nikon Ti2 E800 Scope under a 20X air long range objective, using the Nikon Element software. Images were acquired at the resolution of 2044 pixels x 2048 pixels. With the designated magnification ratio for the objective (1 pixel = 0.23 μm), each field of view has an area of ~160,000 m2. Digital images for each designated color channel were captured at the same exposure time throughout the same plate for quantitative comparison by mean pixel intensity. DAPI images were used to trace the nuclear contour, and bright field images were used to define cell boundaries.

**Quantitative image analysis**

Cell boundaries in the bright field images have sharp and distinguishable local intensity alteration regions (i.e., high-contrast contours, Extended Data Fig. 10a). These high-contrast contours were captured by 2D Gaussian smoothing filter (Matlab built-in image analysis package, with standard deviation of 0.5) and were used to generate cell boundary binary mask by filling up any holes within the contour (Extended Data Fig. 10a). DAPI signals were traced and used to generate binary mask for the nucleus (Extended Data Fig. 10b). Fluorescent intensities of each color channel for a specific target antigen were determined based on the cell or nuclear boundary masks for level and sub-localization.  Specifically, each cell boundary was dilated to 10 and 25 pixels away, and the mean fluorescent intensity of the region between these two dilated boundaries (region between yellow and blue dotted line, Extended Data Fig. 10c,d) was used as local background intensity for a given cell (Green shaded region, Extended Data 10d). The local background intensity was subtracted from the fluorescent pixel intensities within that cell/cell nucleus. This background subtracted value was then summed up and divided by the area (cell or nucleus) of interests to calculate the mean pixel intensity, which was used as present the expression level of target antigens in the cell/cell nucleus. Furthermore, the fluorescence intensity distribution of the negative control dish (treated without the targeted primary antibodies) was used to approximate the "cutoff" fluorescence intensity value for the positive signals (See example in Extended Data Fig. 10e).

To determine the cellular distribution pattern of phosphor-Myosin Light chain (pMLC), the ratio of the relative pMLC level close to the cell edge verses towards the cell center was used. To define the "cell-edge" region, the traced cell boundary (Red solid line, Extended Data Fig. 10d) for each cell was objectively eroded in at a distance that was close to ~1/5 of average cell boundary-to-cell nucleus distance (Black dotted line, Extended Data Fig. 10d). The region within the eroded boundary is thus counted as cell-center region (Brown shaded region, Extended Data Fig. 10d), while the region between the cell boundary and the eroded boundary is considered as cell-edge region.

### Isolation and culturing of muscle satellite cells

All experimental procedures for the mouse were approved by Carnegie Institutional Animal Care and Use Committee (IACUC). Muscle satellite cells were isolated from Pax7-ZSGreen mice[21], and subjected to fluorescence activated cell sorting (FACS) by BD FACSAria III using a protocol modified from Liu et al[52]. Briefly, after euthanasia, hindlimb muscles from 3~6 months mice were dissected out and minced. The minced tissues were put into 10 mL of Wash Medium (10% Horse Serum (HS; Gibco) in Ham's F-10 medium (Thermo Fisher)) containing 1,000 U/mL Collagenase II (Worthington) in a 50 ml conical tube (VWR), and transferred to a shaker water bath set at 37$^{o}$C and 115rpm for 90 minutes of digestion. After digestion, the tube was filled up to 50 mL with cold fresh wash medium, spun down, and the supernatant was aspirated out. 30 mL of wash medium containing 100 U/mL Collagenase II and 1.1 U/mL Dispase (Gibco) was then added into the tube. The mixture was put back into the same shaker water bath for 30 minutes. Afterwards, cell/tissue mixtures were dispersed via aspirating and ejecting through a 20-gauge needle in and out of a 30 mL syringe (Falcon) 10 times. The mixture was then filtered through a 40-μm Nylon cell strainer (Corning) into a fresh 50 mL tube. It was then refilled up to 50 mL with fresh cold Wash Medium, spun down, and the supernatant aspirated out. The cell pellet was resuspended in 1 mL of fresh Wash Medium, transferred into a sorting tube through a 40-μm Nylon cell strainer top (Falcon). The cell-containing solution was diluted by additional 1 mL of fresh Wash Medium, to increase the sorting efficiency.

The ZS-Green$^+$ cells were isolated according to gate settings shown in Fig. 2a., and collected into Wash Medium. To determine sorting efficiency, 150 μl of sorted cell solution was spun onto a glass slide using Cytospin™ centrifuge with cytofunnels and cytoclips (Thermo Scientific). The wetted area (on which the cells were landed) from the glass slide was marked by hydrophobic pen (Vector Laboratories H-400), fixed, permeabilized, blocked, and probed for Pax7 and MyoD expression using the procedures described in the Immunofluorescence and Imaging section. Immunofluorescence and Imaging section.

Isolated cells were seeded in a 24-well plate with plastic bottom (Falcon), unless specified otherwise, at a density of 3,000 to 4,000 cells per well; the well has a diameter of 15.6 mm. The wells were pre-coated with Matrigel (Corning) and Fibronectin (Sigma-Altrich) as follows. Matrigel stock solution was provided by the manufacturer at 13 mg/mL, and Fibronectin stock solution was prepared at 1 mg/mL in PBS. They were diluted into Dulbecco's Modified Eagle's Medium Formulation (DMEM, GIBCO) to a final concentration of 1.3 mg/mL of

Matrigel and 25 μg/mL of Fibronectin as the coating solution. 0.3 mL of coating solution was applied to each well, and incubated in a standard tissue culture incubator (37 $^0$C and 5% $CO_2$) for 1 hour, prior to cell seeding. Seeded cells were cultured in growth medium (with or without compression) under standard conditions (37 $^0$C, 5% $CO_2$) for the duration specified in each set of experiments. The growth medium contains 1% Penicillin-streptomycin (PS, Gibco), 0.01% of Fungin (InvivoGen) and Plasmocin (InvivoGen), 0.1% of chicken extract (M.P. Bimedicals), 10 ng/mL Fibroblast Growth Factor (FGF, BD Bioscience), 5% of HS and 20% of Fetal Bovine serum (FBS, Gibco) in DMEM. To assess the differentiation capacity of compressed cells, they were released from compression and continued to be cultured in growth medium until reaching a density (at 7 days later) similar to that of 5d-U cells (Extended Data Fig. 4). Growth media was then replaced by the differentiation media (10% HS in DMEM) to induce myogenic differentiation for 4 days. For comparative purposes, 5d-U cells were then cultured in differentiation medium for 4 days before they were assayed.

All drug-treatments on the cell culture took place after 2 days of culturing and lasted for 3 more days, as a direct comparison with 2+3d-C and 5d-U cells. The concentration for Y-27632 (Tocris, $K_i$ = 0.14-0.22 and 0.3 μM for inhibiting ROCK1 and 2, respectively) was 25 μM. This was prepared by diluting 100 mM Y-27632 stock solution (in 1X PBS, recommended by the manufacturer as maximum concentration) in growth medium (1:4000 dilution). We chose 25 μM for Y-27632 treatment so that the average decrease of overall pMLC level in 2+3d-Y cells was similar to that of 2+3d-C cells. Similar concentration of 1X PBS was added to the control sets for both 5d-U and 2+3d-C cells (Growth medium + 0.025% 1X PBS). The working concentration for DAPT (Selleckchem.com, IC50 = 20 nM) is 10 μM diluting from 100 μM DAPT stock solution in growth medium with 8% DMSO (1:10 dilution). The working concentration for

DAPT on different types of cells ranges from 0.5 to 100 μM, according to the manufacturer. 10 μM was chosen so that a 40~60% decrease of Notch3 nuclear-to-cytoplasmic ratio has been observed for 5d-U cells. Using higher concentration than 10 μM may result in substantial cell loss after 3-day incubation. We choose the working concentrations of 3.5 and 30 μM for GI254023X (Sigma, IC50 = 5.3 nM) and TAPI-0 (Tocris, IC50 = 50-100 nM) (ADAM 10 and 17 inhibitors) respectively, diluting from stock solution containing 35 μM of GI254023X and 300 μM TAPI in growth medium with 8% DMSO (1:10 dilution). These working concentrations were referenced from the published studies in ref. 52 and 53. We adjusted the working concentrations from the literature so that the resulting Notch3 nuclear-to-cytoplasmic ratio for 5d-U cells was at the similar level compared to that of DAPT-treated 5d-U cells (Fig.4, Main text). Therefore, the differences in Notch activation and cell fate brought by mechanical compression can be directly compared. The control (untreated) sets for both 5d-Ud and 2+3d-C cells were cultured with a growth medium containing 0.8% DMSO from day 2.

**Antibodies used:**

| Target | Primary Antibody | Dilution ratio | Corresponding secondary antibody |
|---|---|---|---|
| Pax7 | Mouse IgG1 (supernatant from hybridoma, DSHB) | 1:5 | Alexa flour Goat anti-mouse IgG1 488 or 647, 1:1000 dilution (Invitrogen) |
| MyoD | Mouse IgG2b (Santa Cruz) | 1:250 | Alexa Flour Goat anti-mouse IgG2b 568 or 647, 1:1000 dilution (Invitrogen) |

| | | | |
|---|---|---|---|
| Notch1 | Rabbit IgG (Abcam) | 1:200 | Alexa Flour Goat anti-rabbit IgG 647, 1:1000 dilution (Invitrogen) |
| Notch3 | Rabbit IgG (Abcam) | 1:200 | Alexa Flour Goat anti-rabbit IgG 647, 1:1000 dilution (Invitrogen) |
| PMLC | Rabbit IgG (Cell Signaling) | 1:100 | Alexa Flour Goat anti-rabbit IgG 647 or 568, 1:1000 dilution (Invitrogen) |
| MF-20 | Mouse IgG2b (supernatant from hybridoma, DSHB) | 1:250 | Alexa flour Goat anti-mouse IgG2b 568 or 647, 1:1000 dilution (Invitrogen) |

**Extended Data Figures and Legends:**

**Extended Data Fig. 1| Cell dimension measurement for in vivo intravital imaging of Pax7-YFP cells, in both cell height (a) and axial length (b).** (Cells are selected from data collected in Ref. 5. 21 cells from uninjured muscle; 11 cells from 1dpi muscle and 9 cells from 3dpi. Data is presented with mean ±s.d. $p$ value was assessed with student's two-tail t-test using Matlab. Comparison was considered significant if $p \leq 0.05$

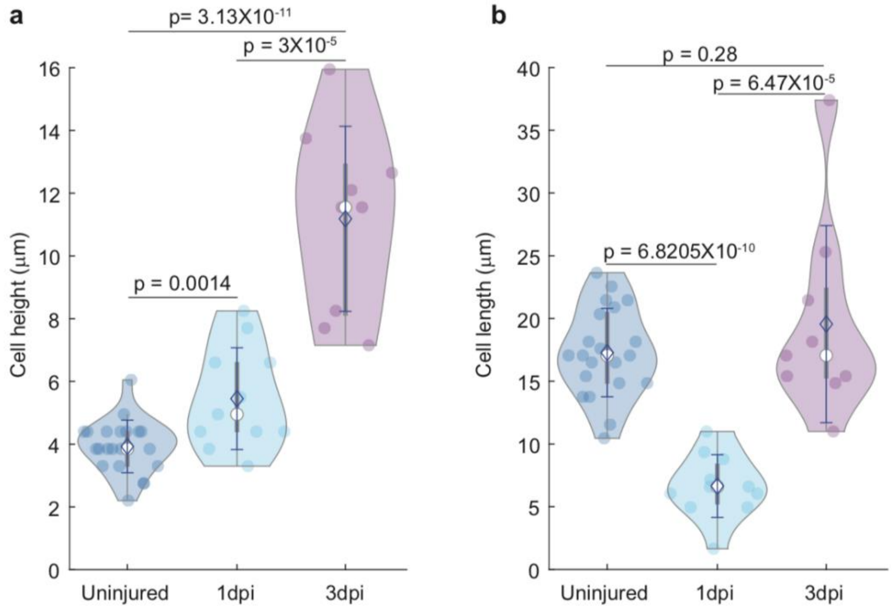


**Extended Data Fig. 2| Illustration and dimensions of compression device. a-b,** Illustration of compression device mold on a silicon wafer **(a),** and designed pillar pattern **(b)**. **c,** Top view of compression pillar. **c,** Height measurement of one pillar on the silicon wafer.

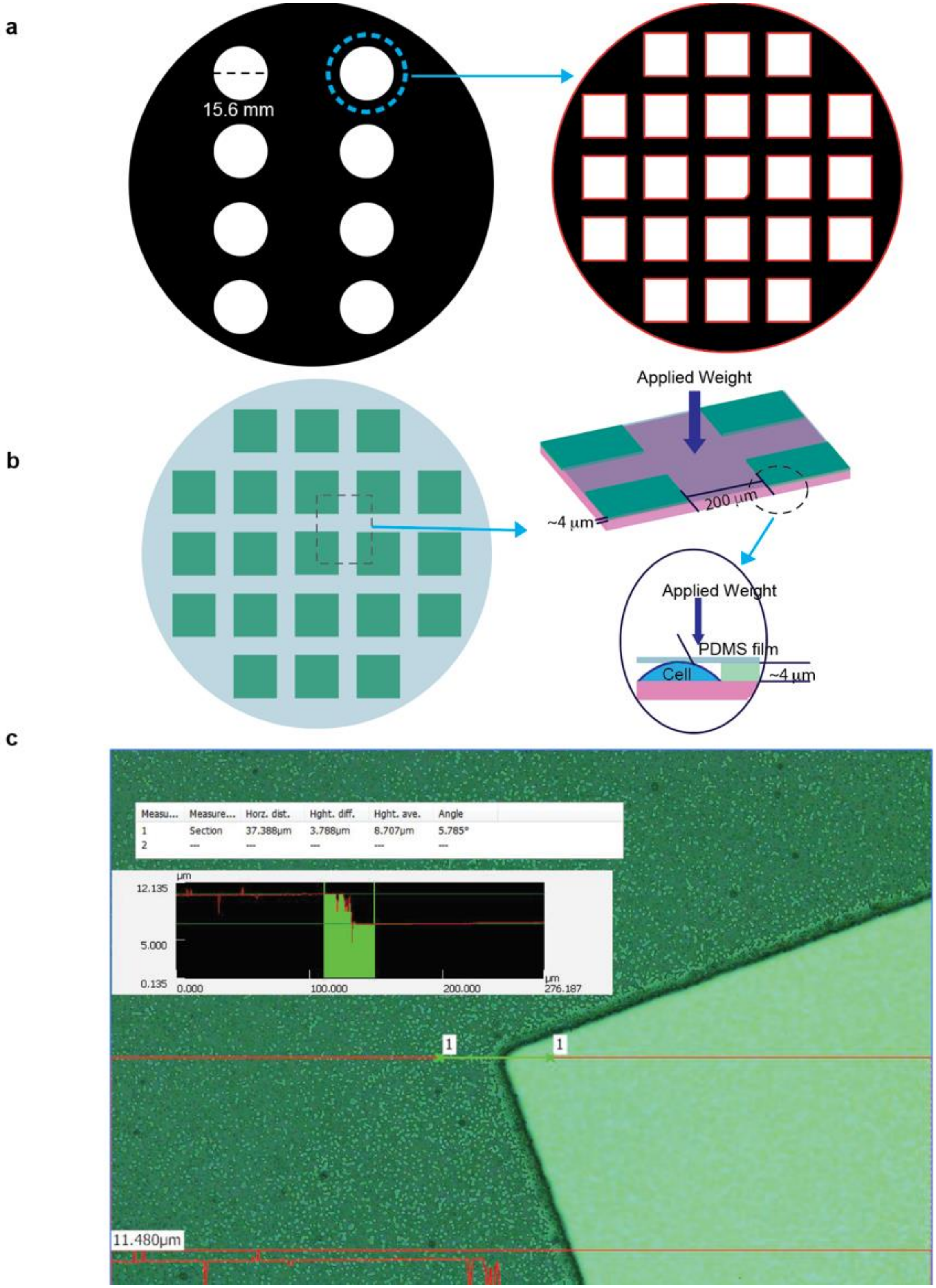
a
15.6 mm
b
Applied Weight
200 µm
~4 µm
Applied Weight
PDMS film
Cell
~4 µm
c
Measu...
Measure...
Horz. dist.
Hght. diff.
Hght. ave.
Angle
1
Section
37.388µm
3.788µm
8.707µm
5.785°
2
µm
12.135
5.000
0.135
0.000
100.000
200.000
276.187
1
1
11.480µm

**Extended Data Fig. 3| 3D confocal measurement of cell height and volume. a,** 3D rendering of uncompressed and compressed cell shape based on confocal data**. b-c,** Measured cell height **(b)** and volume **(c)** for uncompressed and compressed cells. (Four trails for both uncompressed and compressed cells. 19 cells for uncompressed and 28 cells for compressed). Data is presented with mean $\pm$s.d. $p$ value was assessed with student's two-tail t-test using Matlab. Comparison was considered significant if $p \leq 0.05$

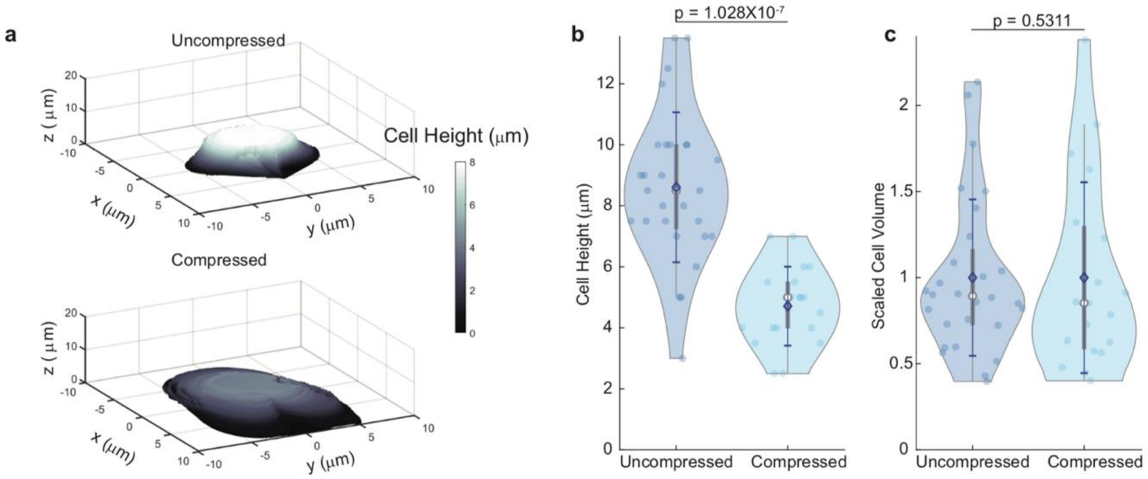


**Extended Data Fig. 4| Compression changes MuSCs' fate. a-c,** Single-channel images for 2d-U **(a)**, 5d-U **(b)** and 2+3d-C **(c)** cells. (Scale bar 25 μm). **(b)** and **(c)** are the same sample presented in **Fig. 1b. d,** Bright-field example of cells right after removing compression pillar (2+3d-C) and after 7-day culturing after compression device's removal (2+3d-C+7) (Scale bar 25 μm). **e,** Cell density comparison between 5d-U and 2+3d-C+7. **f,** Examples 5d-U and 2+3d-C+7 cells after 4-day culturing in differentiation medium, designated as 5d-U+4 and 2+3d-C+7+4, respectively (Scale bar 25 μm). **g,** Examples of 7d-U and 7+3d-C cells (Scale bar 25 μm). **(e)** The sample selected for cell density quantification is the same sample presented in **Fig. 1j,k.**

Data is presented with $\pm$s.d. Ten fields of view were selected for 2+3d-C+7+4 sample and Forty-two fields of view were selected for 5d-U+4 samples. $p$ value was assessed with student's two-tail t-test using Matlab. Comparison was considered significant if $p \leq 0.05$

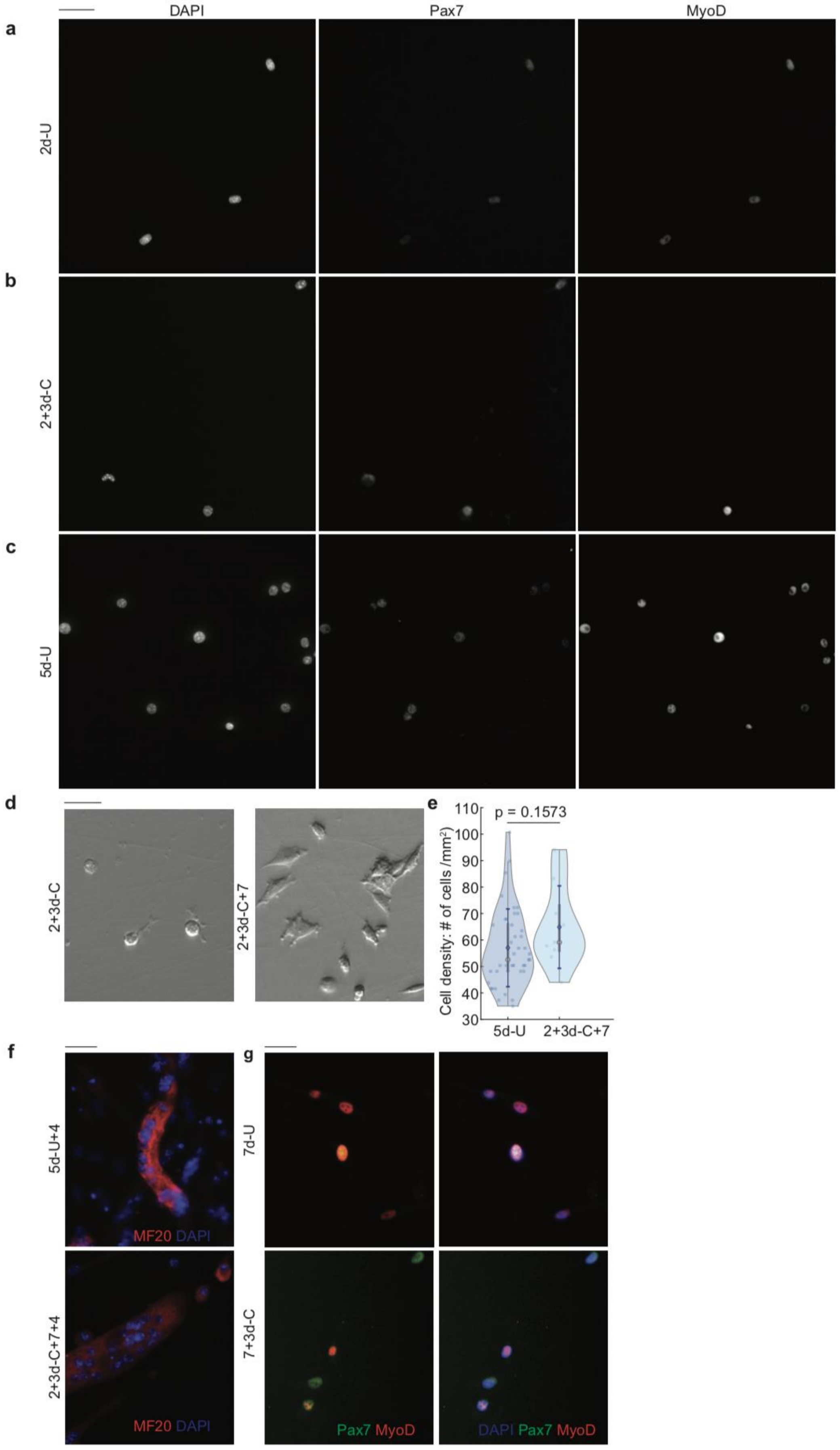
a
b
c
DAPI
Pax7
MyoD
2d-U
2+3d-C
5d-U
d
2+3d-C
2+3d-C+7
e
p = 0.1573
Cell density: # of cells /mm²
110
100
90
80
70
60
50
40
30
5d-U
2+3d-C+7
f
5d-U+4
MF20 DAPI
2+3d-C+7+4
MF20 DAPI
g
7d-U
7+3d-C
Pax7 MyoD
DAPI Pax7 MyoD

**Extended Data Fig. 5| Illustration and coordinate designation of mathematical model.** Cell surface is discretized by azimuthal radius, R and azimuthal angle, θ. This illustration shows uncompressed cell shape (dotted line, with height $H_0$), and predicted cell shape with height, H.

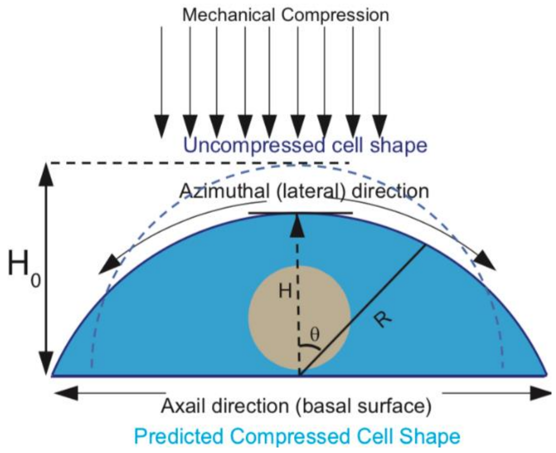


**Extended Data Fig. 6| Manipulating tension by Y-27632 treatment or seeding cells on 12 KPa hydrogel. a,** Examples of pMLC distribution for 5d-U, 2+3d-C and 2+3d-Y cells (Scale bar 25 μm). **b,** Comparison of measured pressure between 5d-U and 2+3d-Y cells by AFM. (5d-U cells pressure data is the same as the plastic data in **Fig. 2g.** Four trails for 2+3d-Y sets, 24 cells). **c,** Cell fate comparison between 5d-U and 2+3d-Y cells. (2+3d-Y's cell fate data is the same as in **Fig. 2e;** Three trails for 5d-U sets, 985 cells). **d,** Cell fate comparisons of 2d-U, 5d-U and 2+3d-C between cells seeded on plastic and 12 KPa hydrogel. (All plastic-seeded cell data is the same as in **Fig. 1c**. All 12 KPa hydrogel-seeded cell data is the same as in **Fig. 2i**). **e,** pMLC Edge/Center ratio comparison for 2+3d-C cells seeded on plastic (>GPa) and 12 KPa hydrogel. (Same data presented in **Fig. 2c** for cells seeded on plastic and **Fig. 2k** for cells seeded on 12 KPa hydrogel). **f,** Single-channel images for 12 KPa hydrogel-seeded cells, both 5d-U and 2+3d-C. Data in **(b)** and (**e**) is presented with $\pm$s.d. $p$ value was assessed with student's two-tail t-test using Matlab. Data in **(c)** and **(d)** is presented by overall fraction. $p$ value was assessed based on two-tailed Cochran-Mantel-Haenszel test using Matlab. Comparison was considered significant if $p \leq 0.05$

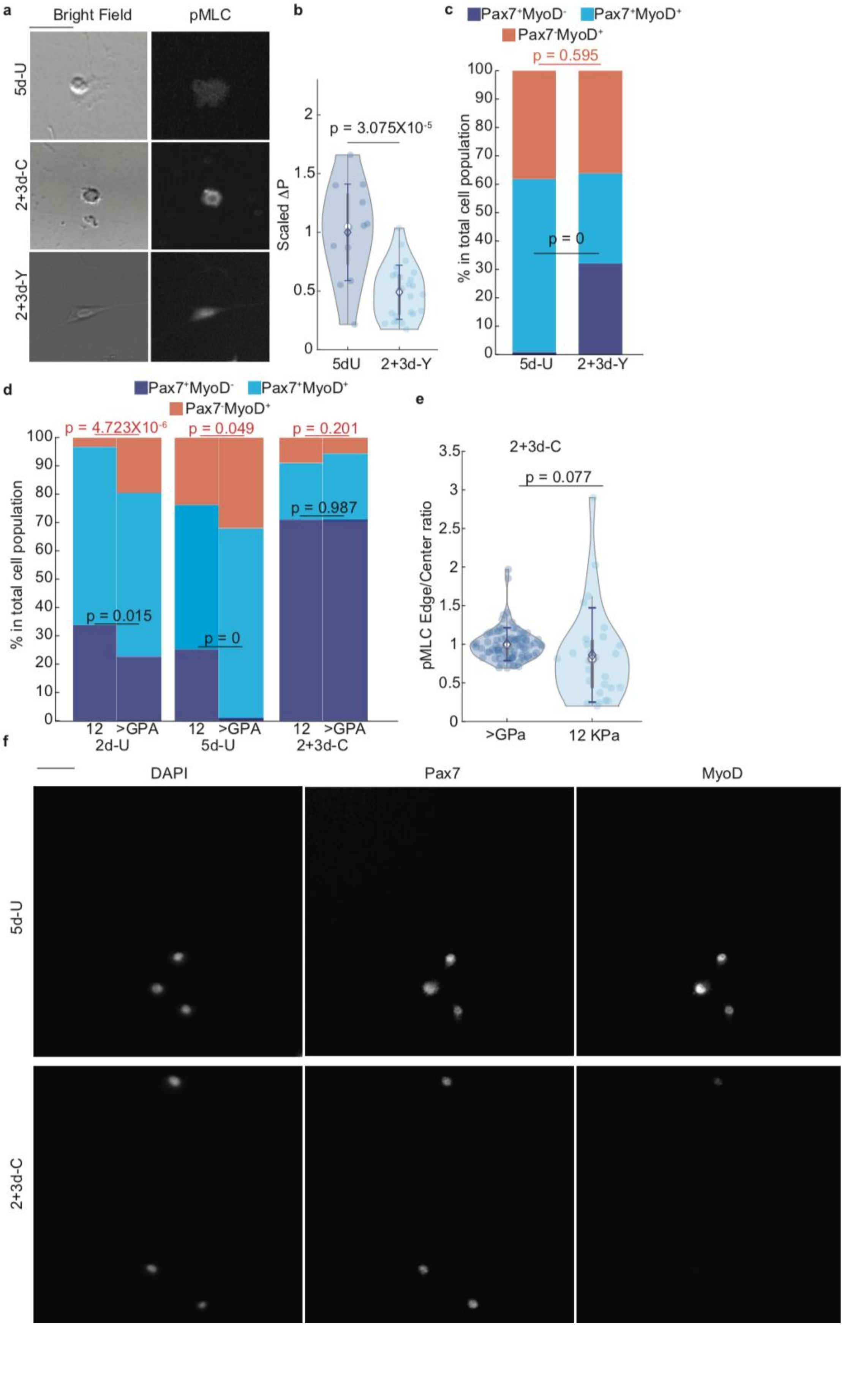

a
Bright Field
pMLC
5d-U
2+3d-C
2+3d-Y
b
p = 3.075X10-5
Scaled ΔP
5dU
2+3d-Y
c
Pax7+MyoD-
Pax7+MyoD+
Pax7-MyoD+
p = 0.595
p = 0
% in total cell population
5d-U
2+3d-Y
d
Pax7+MyoD-
Pax7+MyoD+
Pax7-MyoD+
p = 4.723X10-6
p = 0.049
p = 0.201
p = 0.987
p = 0.015
p = 0
% in total cell population
12
>GPA
2d-U
5d-U
2+3d-C
e
2+3d-C
p = 0.077
pMLC Edge/Center ratio
>GPa
12 KPa
f
DAPI
Pax7
MyoD
5d-U
2+3d-C

**Extended Data Fig. 7| Additional RNA sequence results. a,** PCA plot for 5d-U, freshly isolated (F) and 2+3d-C samples. (PCA scores are generated by R studio). **b,** Comparison of upregulated genes between steady-state mRNA and nascent RNA, based on Ref. 28 and 29, respectively. **c,** Pathway analyses on "core" quiescence genes through reactome, GO (DAVID) and IPA. **d,** Pathway analyses on all upregulated genes in 2+3d-C compared to either 5d-U or freshly isolated cells. $p$ values in **(c)** and **(d)** are calculated from each respective pipeline. The pathway is significantly enriched if $p<0.05$.

**a**

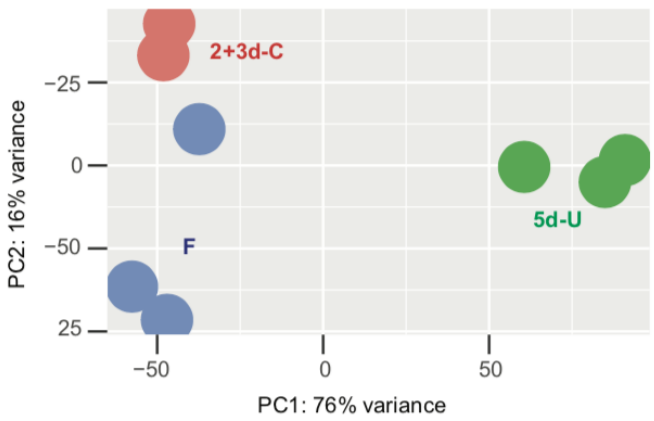


**b**

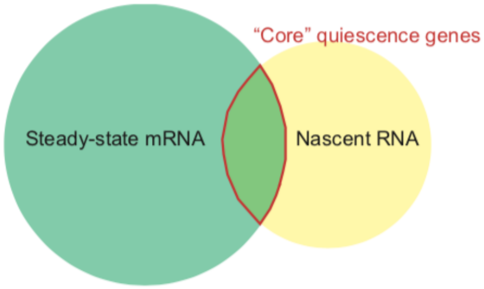


**c**

| Pathway | p value | | |
|---|---|---|---|
| | Reactome | GO (DAVID) | IPA |
| Extracellular Matrix | 5.3X10$^{-4}$ | Adhesion 0.00155 | X |
| Notch signal pathway | Signaling by Notch 0.00155<br>Signaling by Notch1 0.01<br>Signaling by Notch3 0.0121<br>Signaling by Notch4 0.0492<br>NICD1 0.0081<br>NICD3 0.00012<br>NICD4 0.00253 | X | 0.008 |
| GPCR signaling | CAM-PDE1 0.01304 | X | |
| Rho GTPase | 0.043 | X | 0.025 |
| Hedgehog | 0.049 | X | 0.069 |

**d**

| Pathway | p value | |
|---|---|---|
| | GO (DAVID) | IPA |
| GPCR | Response to cAMP 4.5X10$^{-6}$ | GPCR signaling 1.05X10$^{-5}$<br>cAMP-mediated signaling 0.0017 |
| Cell mechanics | Actin cytoskeleton 0.047<br>Rho signal transduction 9.7X10$^{-4}$<br>Calcium ion homeostasis 0.0021<br>Response to mechanical stimulus 3X10$^{-7}$ | Actin cytoskeleton 3.98X10$^{-5}$<br>Rho family GTPases 1.35X10$^{-4}$<br>Calcium transport I 3.63X10$^{-4}$<br>RAC signaling 5.62X10$^{-4}$<br>RHOA signaling 0.0126 |
| WNT pathway | WNT 0.0038<br>Canonical 0.0022 | WNT/Ca+ 0.0017 |
| Notch signaling | 1.5X10$^{-5}$ | 0.0035 |
| Growth factor | X | FGF signaling 0.0155<br>PDGF signaling 0.0054 |
| Cell-ECM | Integrin signaling 0.035<br>Cell adhesion 2.8X10$^{-11}$ | Integrin signaling 0.0062<br>ILK signaling 6.76X10$^{-4}$<br>FAK signaling 0.0051 |

**Extended Data Fig. 8| Examples of 5d-U and 2+3d-C cells.** 5d-U and 2+3d-C bright field and Notch (red)/DAPI (blue) immunofluorescent images for S2-inhibited (+ADAM inhi) **(a)** and S3-inhibited (+γ–s inhi) **(b)** cells (Scale bar, 25 μm).

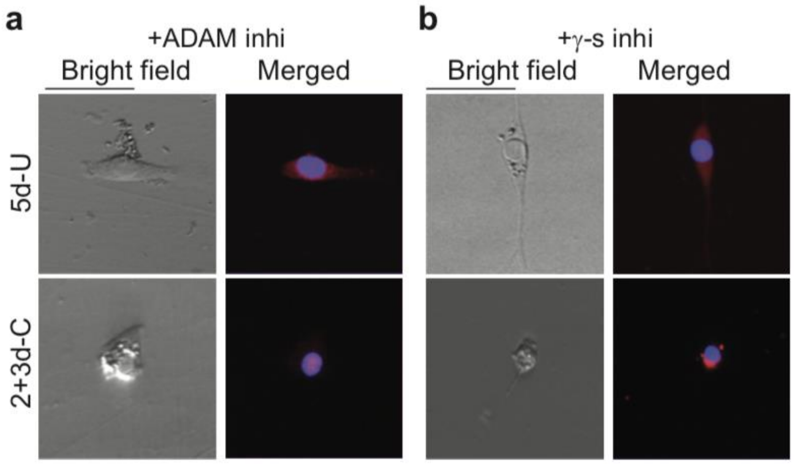


**Extended Data Fig. 9| Technical details of AFM data-fitting. a,** Illustration of AFM cantilever approaching a cell. **b,** Theoretical cell shape before (black) and after (red line) indentation. **c,** Example of indentation strain-responding stress (F) curves of 5d-U cells on plastic and 12 KPa hydrogel, and 2+3d-Y cells. **d-f,** Fitting indentation curves for 5d-U cells on plastic **(d),** and on 12 KPa hydrogel **(e)** and 2+3d-Y cells **(f)** with theoretical indentation strain-$\frac{F}{\Delta P}$ curve to calculate $\Delta P$

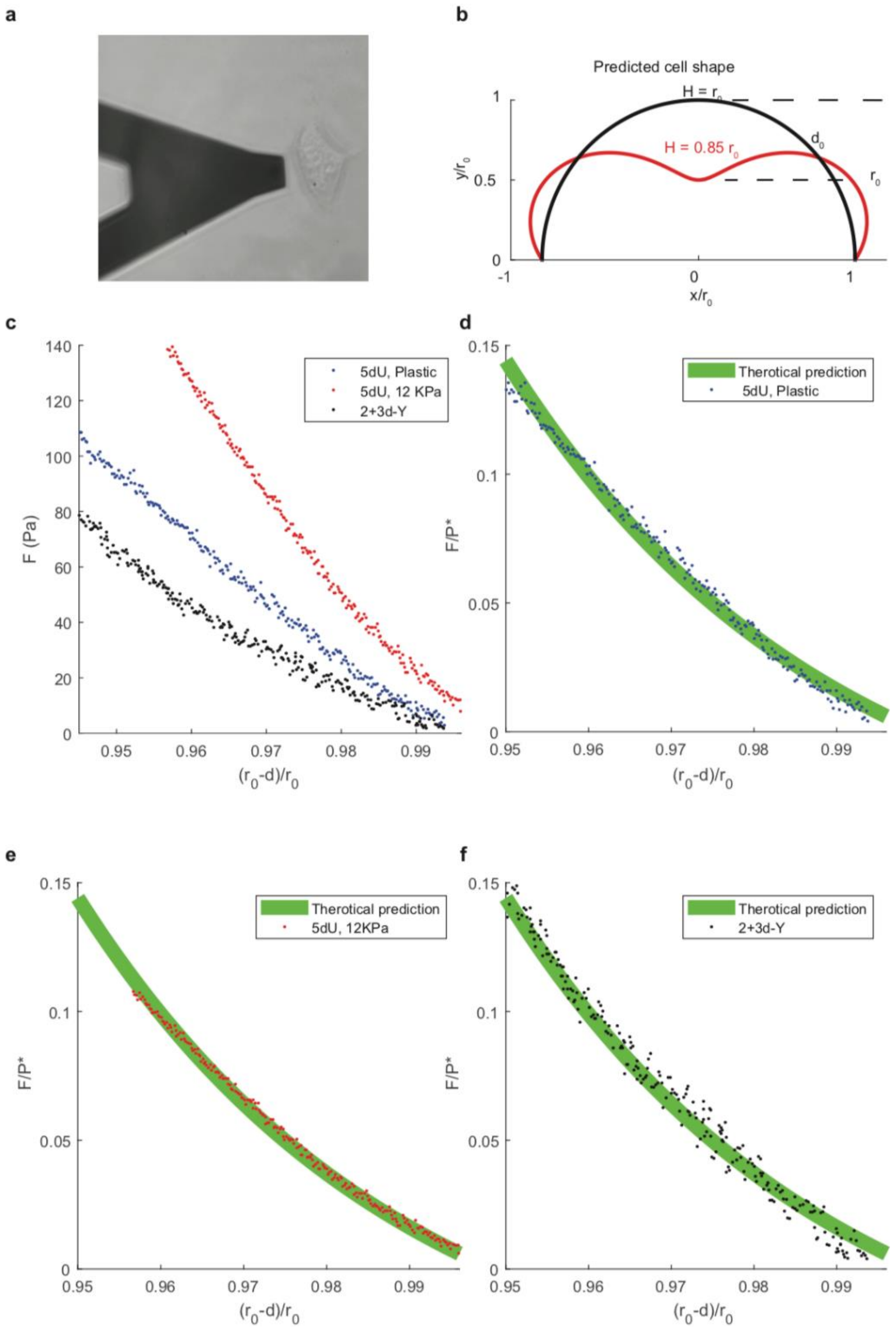


**Extended Data Fig. 10| Quantification for immunostaining data. a,** Example of cell-boundary tracing using bright field. **b,** Example of cell nucleus boundary tracing using DAPI channel. **c-d,** traced cell boundary shown in **(a)** and the respective dilated/eroded boundary. **e,** Example of Pax7 and MyoD intensity histograms for 2+3d-C cells on 12 KPa hydrogel and 5d-U cells on plastic.

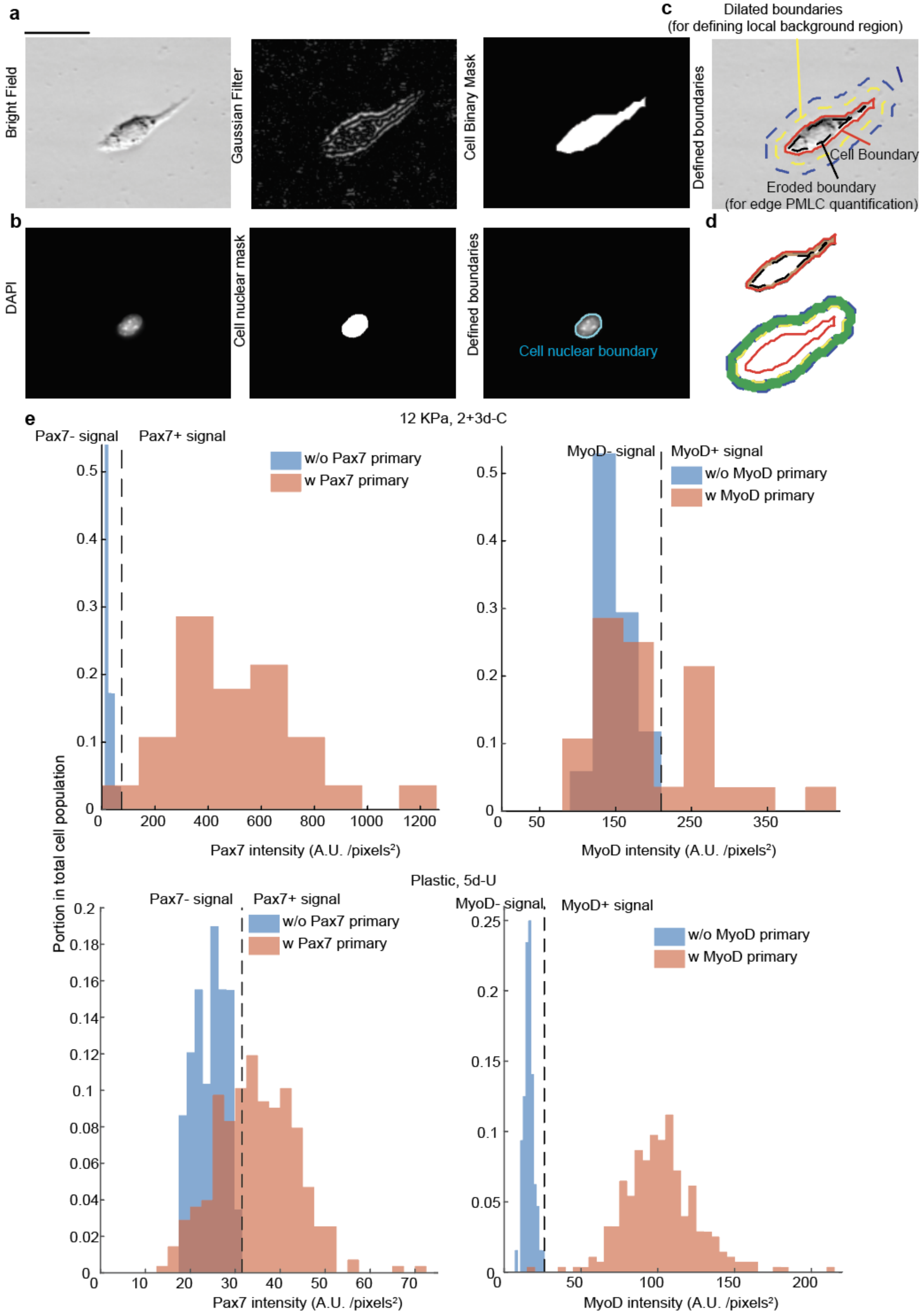
a
Bright Field
Gaussian Filter
Cell Binary Mask
c
Dilated boundaries
(for defining local background region)
Defined boundaries
Cell Boundary
Eroded boundary
(for edge PMLC quantification)
b
DAPI
Cell nuclear mask
Defined boundaries
Cell nuclear boundary
d
e
12 KPa, 2+3d-C
Pax7- signal
Pax7+ signal
w/o Pax7 primary
w Pax7 primary
Pax7 intensity (A.U. /pixels²)
MyoD- signal
MyoD+ signal
w/o MyoD primary
w MyoD primary
MyoD intensity (A.U. /pixels²)
Portion in total cell population
Plastic, 5d-U
Pax7- signal
Pax7+ signal
w/o Pax7 primary
w Pax7 primary
Pax7 intensity (A.U. /pixels²)
MyoD- signal
MyoD+ signal
w/o MyoD primary
w MyoD primary
MyoD intensity (A.U. /pixels²)
